# Zinc isotope fractionation in native and cloned mosses: implications for standardized pollution biomonitoring

**DOI:** 10.64898/2026.10.19.760122

**Authors:** Aridane G. González, Oleg S. Pokrovsky, Álvaro Muñoz-Noval, Jerome Viers, Anna K. Ostendorf, Ralf Reski

## Abstract

Mosses are widely used as biomonitors of atmospheric trace-metal deposition, but metal retention by the moss itself may modify source-related isotope signatures. Here, controlled Zn adsorption experiments, stable Zn isotope measurements, surface acid–base characterization, and Zn K-edge X-ray absorption spectroscopy (XAS) were combined to investigate Zn^2+^ retention and isotope fractionation in four native moss taxa and *in vitro*-cultivated *Sphagnum palustre* clones. Zn^2+^ adsorption was strongly pH-dependent, consistent with progressive deprotonation of reactive surface sites. Moss-associated Zn^2+^ was slightly enriched in heavy isotopes relative to the coexisting solution, with Δ^66^Zn_moss-solution_ generally ranging from +0.01 to +0.09‰, but no systematic relationship was observed between isotopic fractionation and the fraction of Zn^2+^ adsorbed. Linear-combination fitting of X-ray Absorption Near-Edge Structure (XANES) indicated predominantly cellulose-like, O-rich coordination environments, with variable phosphate and minor acetate contributions and only small nominal Zn–S fractions. Extended X-ray Absorption Fine Structure (EXAFS) showed first-shell Zn–O distances of approximately 1.96–2.05 Å and coordination numbers of about 3.6–7.6, confirming substantial structural heterogeneity among samples. Importantly, Δ^66^Zn_moss-solution_ was not correlated with either Zn–O distance or coordination number, indicating that the net isotope effect cannot be described by a single bulk structural parameter. Native and cultivated *Sphagnum* showed comparable isotopic offsets within analytical uncertainty despite differences in surface properties and adsorption capacity. These results indicate that Zn isotope fractionation in mosses reflects the integrated response of multiple binding environments and support the use of standardized *Sphagnum* clones in isotope-based biomonitoring.

## 1. INTRODUCTION

Atmospheric deposition is a major pathway by which trace metals are transferred from emission sources to terrestrial ecosystems, with important consequences for ecosystem functioning and biogeochemical cycling. Direct measurements of wet and bulk deposition provide essential quantitative information, but their spatial coverage is often constrained by the resources and infrastructure required for long-term monitoring. Moss biomonitoring provides a complementary approach that can resolve spatial patterns of atmospheric trace-element exposure and deposition over extended periods. Terrestrial ectohydric mosses are particularly suitable for this purpose because their mineral nutrition is strongly influenced by atmospheric inputs and their tissues efficiently retain deposited trace elements (Harmens et al., 2010; Ares et al., 2012; Fernández et al., 2015).

The suitability of mosses for trace-metal biomonitoring is closely related to their ectohydric physiology and surface chemistry. Because they lack true roots and vascular tissues (Reski, 1998), mosses obtain much of their water and mineral nutrients directly from atmospheric inputs. Their cell walls contain abundant proton- and metal-reactive functional groups, including carboxyl, hydroxyl, and phosphate-containing moieties, which confer a high cation-exchange capacity and promote metal retention through ion exchange and surface complexation (González and Pokrovsky, 2014; Di Palma et al., 2026). The large surface-to-mass ratio and morphological complexity of moss tissues further enhance their adsorption capacity, although substantial interspecific differences exist (Rühling and Tyler, 1971; Aceto et al., 2003; Kłos et al., 2014; González et al., 2016). These characteristics make moss elemental concentrations highly responsive to atmospheric metal loading and underpin the moss-bag technique, in which preconditioned moss material is exposed at selected sites to compare contaminant accumulation under controlled exposure conditions (Ares et al., 2012).

A major limitation of the moss-bag approach, however, is the heterogeneity of field-collected material and the incomplete standardization of experimental protocols. Differences in species, plant age, previous environmental exposure, physiological status, morphology, and sample preparation may influence metal accumulation and complicate comparisons among monitoring sites (Ares et al., 2012; Fernández et al., 2015). The development of *in vitro*-cultivated moss clones therefore offers an attractive route toward standardized biomonitoring material with a controlled exposure history and reproducible physicochemical properties. Within the EU-FP7 MossClone project, a clone of *Sphagnum palustre* was developed and characterized for this purpose. Previous laboratory and field studies demonstrated that cloned *S. palustre* combines a low and stable background elemental composition with high metal-adsorption capacity and enhanced accumulation signals, supporting its use as a standardized material for active biomonitoring (González et al., 2016; Di Palma et al., 2016; Capozzi et al., 2016; Capozzi et al., 2017).

Zinc is particularly relevant in this context because its global biogeochemical cycle has been substantially perturbed by anthropogenic activities, including mining and smelting, metal processing and galvanization, fossil-fuel combustion, traffic-related emissions, and the application of Zn-containing fertilizers. Atmospheric Zn derived from these sources can be transported far from emission areas and contribute substantially to aerosol Zn even in remote environments (Cloquet et al., 2006; Wei et al., 2025; Duan et al., 2025). Beyond total Zn concentrations, stable Zn isotope compositions provide an additional dimension for investigating atmospheric sources and environmental pathways. Although isotopic fields from different sources may partly overlap, variations in Zn isotopes have been successfully used to constrain anthropogenic contributions and Zn cycling in atmospheric and terrestrial systems (Cloquet et al., 2008; Wei et al., 2025; Duan et al., 2025).

The interpretation of Zn isotope signatures is complicated, however, by fractionation occurring during Zn transfer among chemical and biological pools. Experimental studies have demonstrated measurable Zn isotope fractionation during adsorption onto mineral surfaces, complexation with organic matter, and interaction with biological interfaces (Gélabert et al., 2006; Balistrieri et al., 2008; Juillot et al., 2008; Jouvin et al., 2009; Coutaud et al., 2014). Importantly, these effects are related to changes in Zn bonding and speciation. Adsorption onto Fe oxyhydroxides preferentially retains heavier Zn isotopes and has been related to changes in Zn–O bond lengths and coordination relative to aqueous Zn (Balistrieri et al., 2008; Juillot et al., 2008), whereas organic-matter complexation produces fractionation that depends on pH and the affinity of the available binding sites (Jouvin et al., 2009). Consequently, the Zn isotope composition recorded by a moss biomonitor may reflect not only the isotopic signature of atmospheric Zn sources, but also fractionation generated during Zn retention by the moss itself.

At the moss–solution interface, Zn retention is expected to be controlled largely by interactions with reactive groups exposed on the cell wall. Carboxyl, phosphate-containing, and other oxygen-bearing functional groups provide important metal-binding sites, while their abundance, acid–base properties, and accessibility determine the net surface charge and adsorption behavior (González and Pokrovsky, 2014). Morphological properties such as surface development may further affect the number and accessibility of these sites, as demonstrated by the substantial differences in Zn adsorption capacity among moss species and between field-grown and cloned *Sphagnum* (Kłos et al., 2014; González et al., 2016). Therefore, differences in surface chemistry and morphology may influence not only the amount of Zn retained but also its local coordination environment and, potentially, the associated isotope fractionation.

Resolving these mechanisms requires structural information beyond bulk Zn concentrations. X-ray absorption spectroscopy (XAS), including X-ray absorption near-edge structure (XANES) and extended X-ray absorption fine structure (EXAFS), provides element-specific constraints on Zn speciation and local coordination in complex biological matrices. Previous spectroscopic studies illustrate the diversity of Zn-binding environments at biological interfaces. On diatom surfaces, Zn is predominantly tetrahedrally coordinated by oxygen and bound to carboxylate ligands (Pokrovsky et al., 2005), while EXAFS investigations of bacterial and fungal systems have identified variable contributions from carboxyl, phosphate, and sulfhydryl ligands depending on metal loading, organism, and physicochemical conditions (Guiné et al., 2006; Fomina et al., 2006, 2007). These observations demonstrate that total Zn accumulation alone cannot predict its molecular-scale binding environment.

Combining Zn isotope measurements with XAS-derived structural information therefore provides a means of testing whether isotope fractionation is associated with particular Zn species, ligand environments, coordination numbers, or Zn–ligand distances. This relationship is especially relevant because previous experimental studies have linked Zn isotope fractionation to changes in coordination and bonding during adsorption and organic complexation (Balistrieri et al., 2008; Jouvin et al., 2009). However, the extent to which such molecular-scale controls operate during Zn adsorption by mosses remains largely unclear. In particular, it is not established whether variations in the magnitude of the isotopic offset between moss and solution are systematically related to the surface chemistry of the moss or to the local structural environment of adsorbed Zn. Resolving this uncertainty is important for evaluating whether moss biomonitors preserve atmospheric Zn isotope signatures or modify them during metal accumulation.

In this study, we combined controlled Zn^2+^ adsorption experiments, stable Zn isotope measurements, surface acid–base characterization, and Zn K-edge X-ray absorption spectroscopy to investigate the mechanisms governing Zn retention and isotope fractionation in native and *in vitro*-cultivated mosses. We tested the hypothesis that Zn isotope fractionation during adsorption is associated with the chemical properties of the moss surface and the local coordination environment of retained Zn. Specifically, we aimed to (i) quantify Zn adsorption and the associated isotopic offset between moss biomass and solution; (ii) determine Zn speciation and local coordination using XANES and EXAFS; (iii) assess the extent to which native and cloned moss materials exhibit comparable Zn-adsorption and isotope-fractionation behavior; and (iv) test whether covaries with spectroscopic and surface-chemical parameters. By directly linking Zn isotope fractionation with molecular-scale binding properties, this study provides a mechanistic basis for evaluating the extent to which moss biomonitors preserve or modify atmospheric Zn isotope signatures during metal accumulation.

## 2. MATERIALS AND METHODS

### 2.1. Chemicals

Analytical-grade reagents and ultrapure water (18.2 MΩ·cm) were used throughout. Zinc nitrate supplied Zn^2+^, with 0.01 M NaNO₃ as background electrolyte. MES and HEPES controlled pH at 5.5 and 6.5, respectively. Reagent preparation and cleaning procedures are detailed in Supplementary Text S1.1.

### 2.2. Adsorption experiments

Intact moss tissues, rinsed three times with ultrapure water and devitalized at 120 °C overnight, were used at 1.0 g dry weight L^−1^. Suspensions were gently stirred and purged with N₂. The pH-edge experiments covered pH 1.8–9.9 at an initial Zn concentration of 3.3 mg L^−1^ (51 µmol L^−1^); adsorption isotherms covered 4–3500 µmol L^−1^ at approximately pH 6.5. Equilibration lasted at least 10 min after each pH adjustment and 10 min after each Zn addition, based on preliminary kinetics. Suspensions were filtered through 0.45-µm cellulose acetate membranes and filtrates acidified for Zn analysis. Procedural blanks and moss-leachate controls were processed in parallel (Supplementary Text S1.2).

### 2.3. Moss samples

*Hypnum* sp., *Sphagnum denticulatum*, *Pseudoscleropodium purum* and *Brachythecium rutabulum* were collected from non-urban Galicia (NW Spain), away from direct pollution sources (González and Pokrovsky, 2014). *Sphagnum palustre* clones 2a and 12a were produced at the University of Freiburg within the EU-FP7 MossClone project (Beike et al., 2015; González et al., 2016; Heck et al., 2021a,b). Surface acid–base properties were characterized by potentiometric titration and Linear Programming Model analysis following González and Pokrovsky (2014) and González et al. (2016) (Supplementary Text S1.3)

### 2.4. Zn concentration and stable isotopic fractionation

Dissolved Zn was measured by inductively coupled plasma mass spectrometry (ICP-MS) and atomic absorption spectrometry (AAS), with quantification limits of 0.6 and 35 µg L^−1^, respectively. Mosses were acid-digested following Gélabert et al. (2006) and Viers et al. (2013), and Zn was purified by anion-exchange chromatography (Maréchal et al., 1999); only samples with 95–100% Zn recovery were analyzed isotopically. In total, 68 solid- and aqueous-phase samples were analyzed using Neptune and P54 multicollector ICP-MS instruments following Coutaud et al. (2014) and Viers et al. (2015). Isotope compositions are expressed relative to JMC batch 3-0749 L:

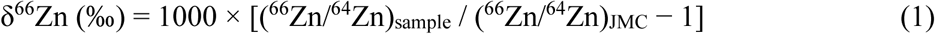

Isotope values are means of 3–5 measurements; reported uncertainties are standard errors, with a minimum of 0.008‰ derived from seven biofilm-standard measurements. Detailed digestion, purification, instrumental and quality-control procedures are provided in Supplementary **Text S1.4**.

### 2.5. X-ray absorption spectroscopy

Zn K-edge X-ray absorption spectroscopy (XAS) was performed at BM25-SpLine, ESRF (Grenoble, France), in fluorescence mode at approximately 10 K. After adsorption, mosses were briefly rinsed, blotted, frozen in liquid nitrogen and stored at −80 °C. X-ray absorption near-edge structure (XANES) spectra were analyzed by least-squares linear-combination fitting of measured Zn references in ATHENA. Extended X-ray absorption fine structure (EXAFS) parameters were fitted using FEFF-calculated Zn–O and, where indicated by XANES, Zn–S paths in ARTEMIS. Acquisition, reference compounds and processing are detailed in Supplementary Text S1.5.

## 3. RESULTS

### 3.1. Zinc adsorption onto mosses

The surface acid–base properties of the moss materials used in this study have been characterized in detail in our previous work (González and Pokrovsky, 2014; González et al., 2016). Zn^2+^ adsorption exhibited a pronounced pH dependence for all four native moss species (**Fig. 1**). At an initial Zn^2+^ concentration of 3.3 mg L⁻¹, adsorption increased from 0–2.7% at pH 2.04–2.33 to 48.6–62.2% at pH 7.14–7.20 (**Table S1**).

**Figure 1.**
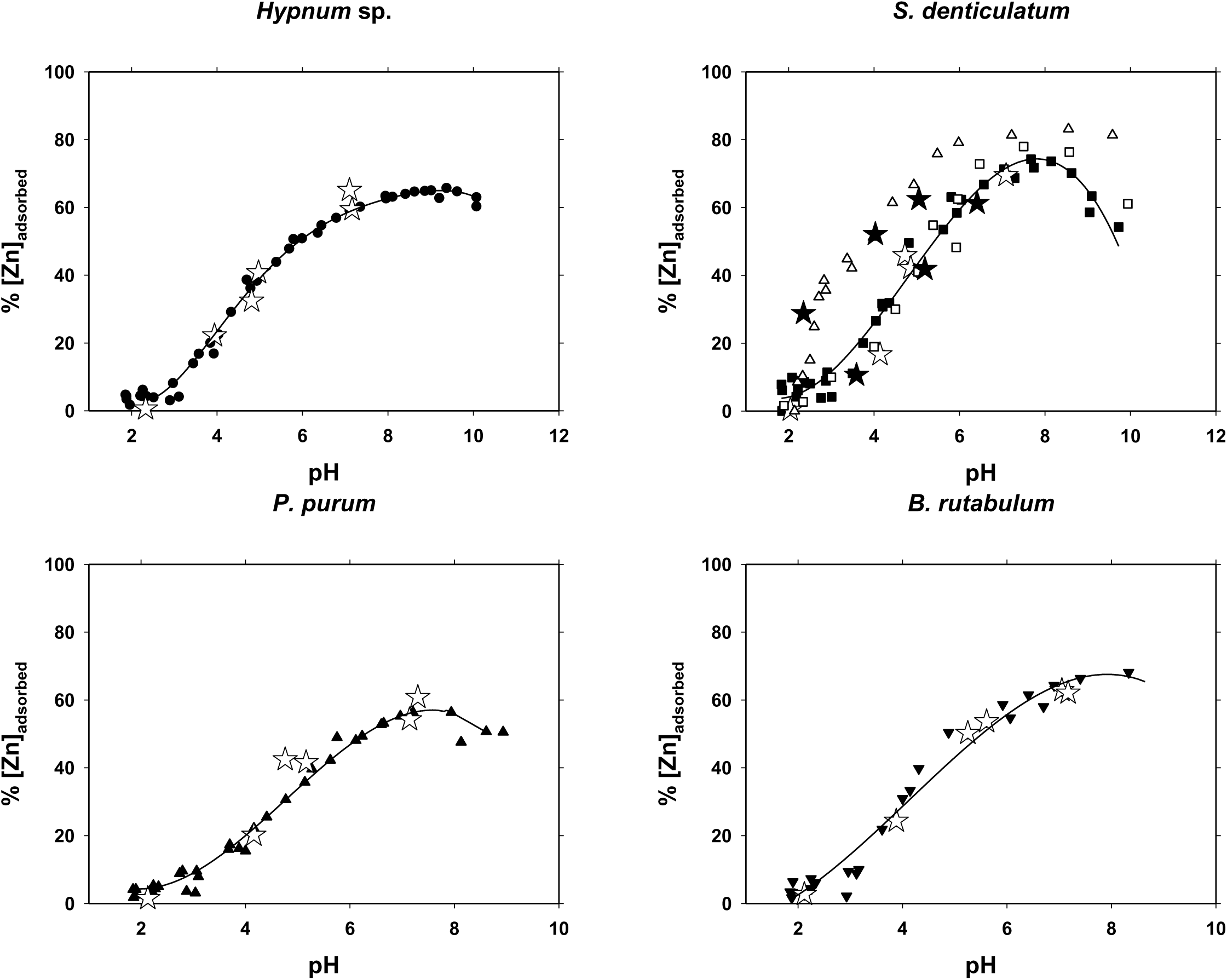
pH dependence of Zn^2+^ adsorption by native and *in vitro*-cultivated mosses. Adsorption experiments were performed at an initial Zn concentration of 3.3 mg L^−1^ (51 µmol L^−1^), a moss biomass concentration of 1.0 g dry weight L^−1^, and an ionic strength of 0.01 M NaNO^3^. Filled black symbols represent native moss samples; open triangles represent the *in vitro*-cultivated *Sphagnum palustre* clone-2a, and open squares represent clone-12a. Stars indicate samples selected for Zn isotope fractionation analysis. Solid lines represent the LPM fitting.

At approximately circumneutral pH, differences among the native moss species were comparatively small. Long-term experiments with *Sphagnum* showed decreasing fractional Zn^2+^ retention with increasing metal loading (**Table S1**). Although native and cloned *Sphagnum* share broadly comparable acid–base behavior, the cloned material provides a greater density of accessible binding sites and a higher Zn^2+^ adsorption capacity on a dry-mass basis (González et al., 2016). Detailed surface properties and adsorption results are provided in the text accompanying **Table S1**.

### 3.2. Stable isotopic fractionation of Zn on mosses

Zn isotope compositions of the moss biomass and corresponding aqueous solutions are reported in Table S2 and illustrated in **Fig. 2**. In most experiments, the Zn associated with moss biomass was isotopically heavier than the coexisting solution, resulting in positive moss–solution isotope offsets (Δ^66^Zn_moss-solution_). The magnitude of this isotopic offset was small, generally ranging from approximately 0.01 to 0.09‰ across the investigated moss species and experimental conditions. No systematic relationship between Δ^66^Zn_moss-solution_ and the fraction of Zn^2+^ adsorbed was apparent across the four native moss species (**Fig. 3a**). Across the investigated adsorption range, the isotopic compositions measured for the cloned material overlapped with those obtained for native *Sphagnum* within analytical uncertainty (**Fig. 3b**). No systematic difference between *S. palustre* clones 2a and 12a was apparent. Isotope definitions, species-specific values and the clonal comparison are detailed in the text accompanying **Table S2**.

**Figure 2.**
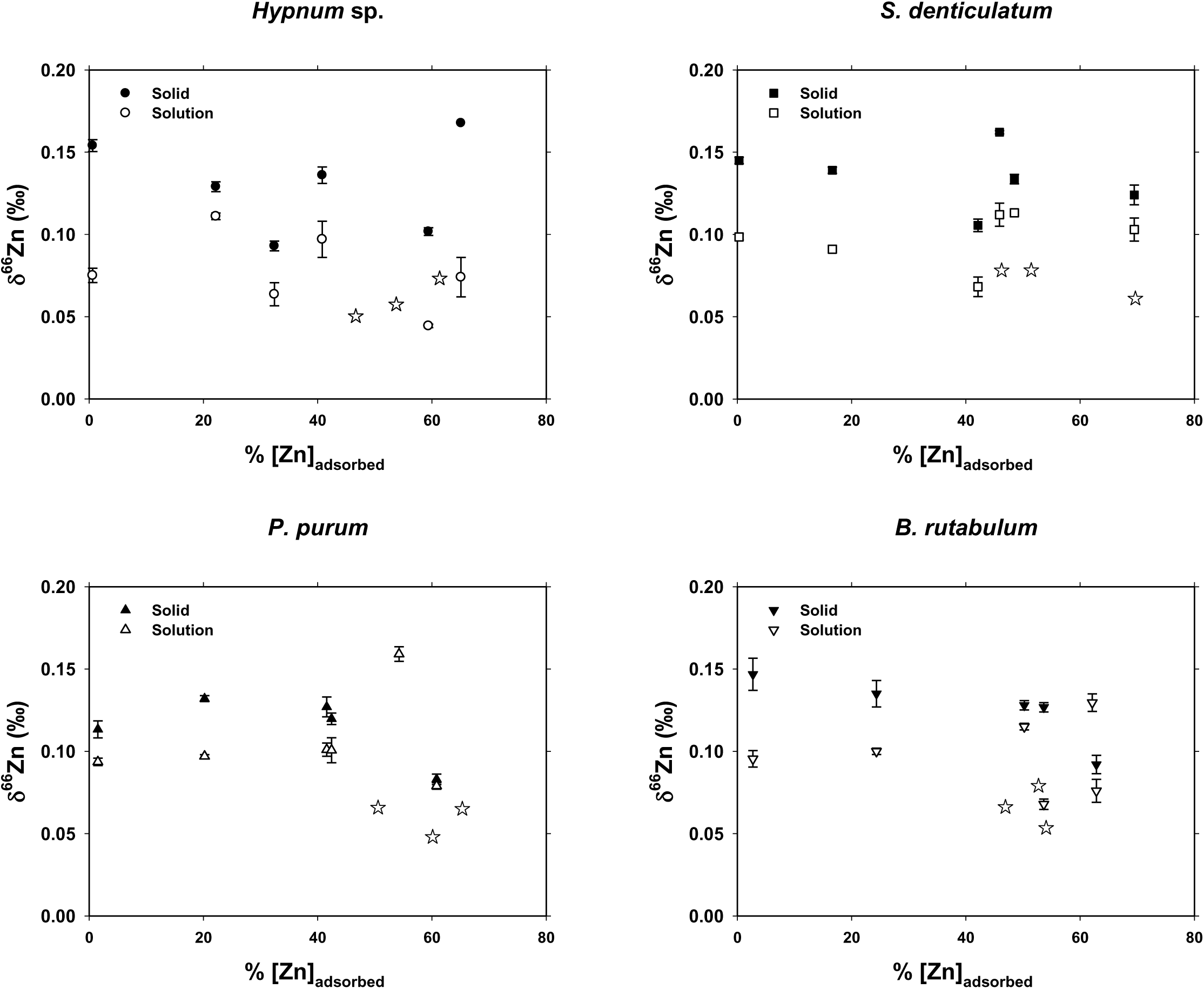
Zn isotope composition of moss-associated and dissolved Zn as a function of the percentage of Zn adsorbed by native mosses. Adsorption experiments were performed at a moss biomass concentration of 1.0 g dry weight L^−1^ and an ionic strength of 0.01 M NaNO_3_. Filled black symbols represent δ^66^Zn values of Zn associated with the solid moss biomass, whereas open symbols represent δ^66^Zn values of the corresponding aqueous solutions. Stars indicate experiments performed at an initial Zn concentration of 0.03 mg L^−1^ (0.46 µmol L^−1^). Error bars represent the analytical uncertainty of the isotope measurements.

**Figure 3.**
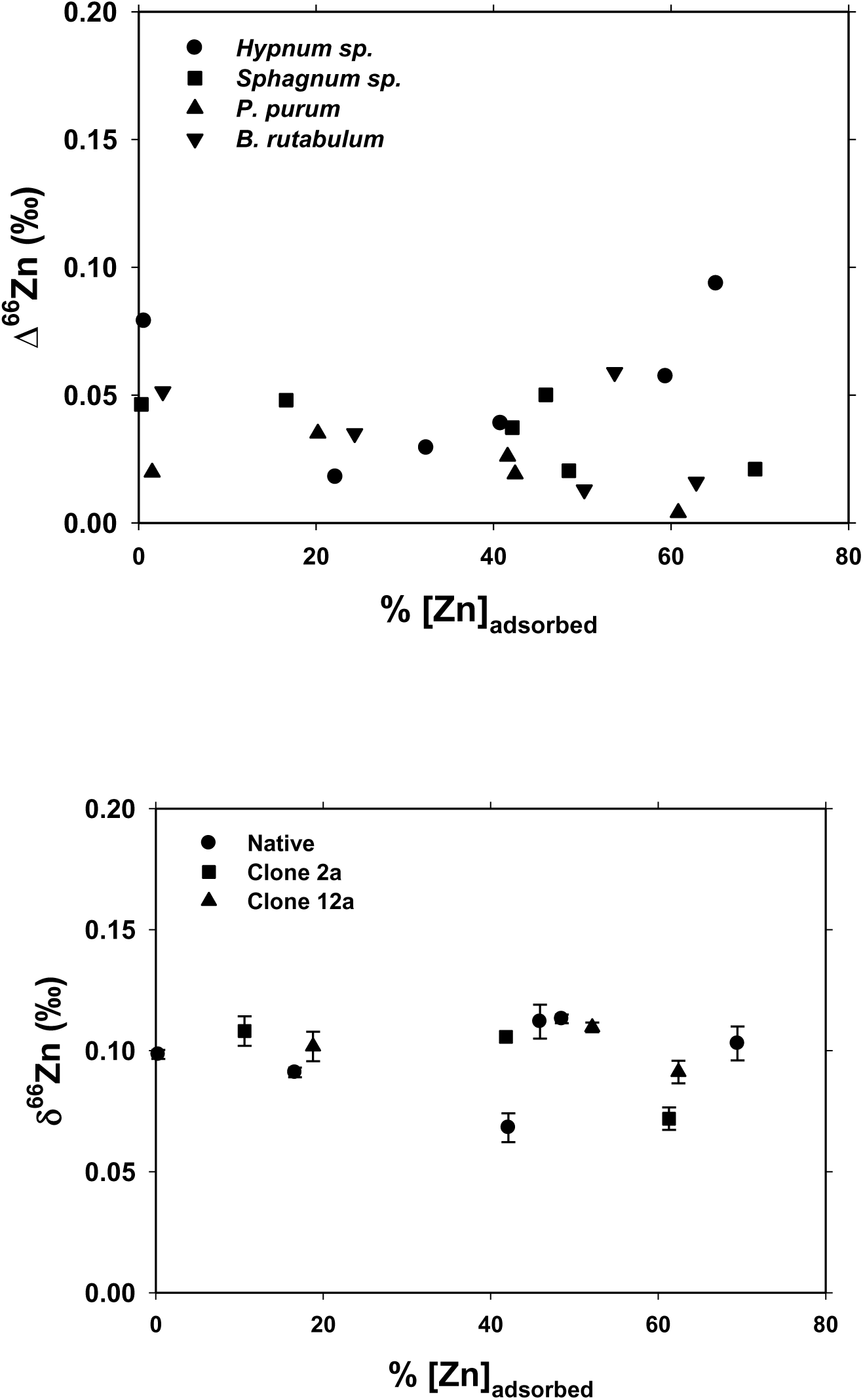
Relationship between Zn^2+^ adsorption and Zn isotope fractionation in native and *in vitro*-cultivated mosses. (a) Zn isotopic offset between moss biomass and the coexisting solution (Δ^66^Zn_moss-solution_) as a function of the percentage of Zn^2+^ adsorbed for the four native moss species: *Hypnum* sp. (circles), *Sphagnum denticulatum* (squares), *Pseudoscleropodium purum* (upward triangles), and *Brachythecium rutabulum* (downward triangles). (b) δ^66^Zn of moss-associated Zn as a function of the percentage of Zn adsorbed for native *Sphagnum* (circles) and the *in vitro*-cultivated *Sphagnum palustre* clones-2a (squares) and -12a (triangles). Error bars represent the analytical uncertainty of the isotope measurements.

### 3.3. XANES analysis

Reference spectra, experimental spectra and their linear-combination fits, and absorption-edge shifts are presented in **Fig. S1**, **Fig. S2** and **Fig. S3**, respectively. Linear-combination fitting of the moss XANES spectra showed that cellulose-like coordination environments represented the predominant Zn component in most samples (**Table S3**). Overall, acetate was detected only as a minor component in most moss samples and rarely exceeded approximately 10–12%, whereas phosphate contributions varied much more strongly among experimental series. Phosphate contributions reached approximately 40–52% in selected samples. Small nominal ZnS fractions, generally below 5%, were below the approximate resolution of the XANES fitting and should be considered indicative rather than quantitatively robust. Series-specific contributions and fitting limitations are detailed in the text accompanying **Table S3** and **Fig. S2**.

### 3.4. EXAFS analysis

EXAFS spectra and fits are shown in Fig. S4, with first-shell structural parameters in Table S4. For most samples, the first coordination shell was adequately described by Zn–O scattering paths. Excluding the atypical Zn-4 fit, Zn–O bond distances ranged from approximately 1.96 to 2.05 Å, while fitted coordination numbers varied from about 3.6 to 7.6. Minor Zn–S coordination was required in only a subset of samples, in agreement with the general predominance of O-bearing coordination environments identified by XANES. Series-specific structural results, Zn–S contributions are provided in the text accompanying **Table S4**.

To examine whether the small Zn isotope fractionation observed during moss adsorption was related to the local coordination environment of retained Zn, Δ^66^Zn_moss-solution_ was compared with the XAS-derived structural parameters for samples for which paired isotope and spectroscopic measurements were available. No significant monotonic relationship was observed between Δ^66^Zn_moss-solution_ and either the first-shell Zn–O distance (R; Spearman ρ = −0.04, *p* = 0.915, *n* = 9) or coordination number (N; ρ = 0.14, *p* = 0.723, *n* = 9). The Debye–Waller factor also showed no statistically significant relationship with the isotopic offset (ρ = 0.55, *p* = 0.125). XANES-derived contributions from cellulose-like and acetate-associated Zn displayed weak trends with Δ^66^Zn_moss-solution_, but these relationships were not significant using rank correlation (|ρ| = 0.61, *p* = 0.079). Evaluation of ZnS and phosphate-related contributions was not warranted for the paired dataset because these components showed little or no variation among the samples for which isotope measurements were available. Overall, the magnitude of Zn isotope fractionation was not systematically related to the principal first-shell structural parameters resolved by EXAFS.

## 4. DISCUSSION

Interpreting Zn isotope fractionation during adsorption onto and uptake by mosses requires identifying the dominant adsorption reactions and linking them to potential Zn-binding surface functional groups, using constraints from in situ structural analysis. We first relate macroscopic adsorption to surface-site properties, then use XAS to characterize the retained Zn environments. These constraints provide the basis for interpreting isotope fractionation, testing structural explanations and defining the opportunities and limits of standardized moss biomonitoring.

### 4.1. Surface-site availability controls Zn retention and adsorption capacity

The strong pH dependence of Zn adsorption observed here is consistent with the amphoteric properties of moss cell walls and their measured pH_PZC_ values (4.6–6.2), defined operationally by zero net proton adsorption (González and Pokrovsky, 2014; González et al., 2016). Proton competition limits Zn binding at low pH, whereas deprotonation of carboxyl and other O-bearing surface moieties increases the availability of negatively charged sites toward circumneutral conditions. The LPM-derived distribution of apparent p*K*_a_ values resolves five tentative site classes, assigned to phosphodiester, carboxyl, phosphoryl, amine, and polyphenolic functionalities, indicating substantial chemical heterogeneity at the moss–solution interface. This diversity is consistent with cell-wall polysaccharides, uronic acids, phenolic compounds and other functionalized macromolecules capable of binding both protons and metals (Clymo, 1963; Popper and Fry, 2003). The adsorption edge can therefore be interpreted as progressive activation of diverse sites rather than the operation of a single adsorption reaction. Broadly comparable pH-dependent adsorption patterns were previously reported for *Hypnum* sp., *Sphagnum* sp., *P. purum* and *B. rutabulum*, despite quantitative species differences (González and Pokrovsky, 2014).

Zn removal reached a steady state within approximately 10 min and remained essentially unchanged during prolonged exposure, supporting predominantly surface-controlled retention. This agrees with rapid cation exchange in *Sphagnum* cell walls (Clymo, 1963) and rapid Zn and Cd adsorption by moss biomass (Kłos et al., 2014). Because the mosses used in the adsorption experiments were devitalized, metabolically controlled intracellular uptake should have been strongly suppressed. Ion exchange and surface complexation at pre-existing cell-wall sites are therefore the most plausible dominant mechanisms under these conditions, although diffusion within the biomass may also contribute to the retention kinetics.

The declining percentage of Zn adsorbed with increasing loading is consistent with progressive occupation of a finite, heterogeneous binding-site population. High fractional retention at low concentrations is compatible with abundant available high-affinity sites relative to dissolved Zn; their progressive occupation may increase the contribution of lower-affinity sites at higher loading. This behavior agrees with adsorption isotherms previously obtained for the same moss species, which showed broadly comparable Zn adsorption but quantitative differences in adsorption capacity (González and Pokrovsky, 2014). Independent moss adsorption experiments similarly related species-dependent differences, at least partly, to surface chemistry, surface development and morphology (Kłos et al., 2014).

The comparison between field-grown and *in vitro*-cultivated *Sphagnum* further suggests that metal retention depends not only on the chemical nature of the surface functional groups but also on their abundance and accessibility. González et al. (2016) showed that cultivated *S. palustre* clones possess larger specific surface areas, higher mass-*n*ormalized densities of proton-reactive sites, and greater Zn adsorption capacities than field-grown *Sphagnum*, while retaining broadly comparable acid–base characteristics. These differences can increase adsorption capacity without requiring a fundamentally different binding mechanism or greater intrinsic affinity at individual sites. Recent comparisons of other *Sphagnum* species with their conspecific clones likewise found broadly conserved surface chemistry and reduced interspecific variability under clonal cultivation, but not a universal increase in binding-site density (Di Palma et al., 2026). The enhanced capacity of the *S. palustre* clones should therefore be regarded as a property of the materials compared here rather than a general consequence of cloning. For biomonitoring, reproducible material production and quantitative equivalence are distinct: identical atmospheric loading need not yield identical dry-mass Zn concentrations in native and cloned mosses. Their accumulation signals should therefore be compared using material-specific calibration and standardized pretreatment and exposure conditions.

### 4.2. Heterogeneous O-donor environments dominate Zn coordination

XANES and EXAFS indicate that moss-associated Zn occupies chemically heterogeneous, predominantly O-coordinated environments, consistent with the acid–base titration results. The dominant Zn-cellulose XANES component reflects an O-rich polysaccharide-like environment rather than demonstrating direct or exclusive binding to cellulose hydroxyl groups. Acetate-associated contributions were minor, whereas phosphate-associated fractions were greater in selected adsorption-isotherm, intact-biomass and growth-clone samples than in most pH-edge samples, reaching approximately 23–52% in the analyzed growth-clone samples (Zn-47, Zn-49 and Zn-51). Such diversity is consistent with the importance of carboxylate and phosphate ligands reported for diatoms (Pokrovsky et al., 2005), bacteria (Guiné et al., 2006) and fungi (Fomina et al., 2007). The phosphate component indicates phosphate-like local coordination but does not, by itself, establish or exclude discrete Zn-phosphate precipitation. Similarly, the absence of major ZnO, ZnCO_3_, Zn citrate, Zn_2_SiO_4_ or metallic Zn contributions from the fits supports predominantly biomass-associated retention within the available reference set, but does not exclude minor components below the approximate 10% practical resolution of XANES fitting.

EXAFS yielded Zn–O distances mostly within approximately 1.96–2.05 Å and fitted coordination numbers of 3.6–7.6. Growth clones and intact biomass generally exhibited lower coordination numbers, close to four, whereas several pH-edge samples approached six to eight. These differences suggest changes in the relative abundance and structure of Zn-binding environments rather than a single structural transformation. Because fitted coordination numbers represent population averages affected by structural disorder and covariance among fitting parameters, they do not uniquely establish molecular geometries. The measured distances are nevertheless compatible with organically coordinated Zn in other biological matrices (Pokrovsky et al., 2005; Guiné et al., 2006; Fomina et al., 2007).

Possible minor Zn–S coordination was indicated by improved EXAFS fits incorporating Zn–S paths in some samples and small nominal XANES ZnS contributions. The latter were generally below 5%, beneath the approximate quantitative resolution, and therefore do not establish an abundant Zn–S-bound fraction. Sulfhydryl ligands identified during bacterial Zn adsorption provide a plausible biological analogue (Guiné et al., 2006). The moss Zn–S signal could likewise reflect thiol or other reduced-S groups, although their molecular identity cannot be resolved by the present data.

### 4.3. Zn retention produces small isotope offsets relative to many sorption systems

The predominantly positive Δ^66^Zn_moss–solution_ values observed in the present experiments suggest slight preferential retention of heavier Zn isotopes by moss biomass. The magnitude of fractionation was small, generally between +0.01 and +0.09‰, and individual offsets near zero should be interpreted within their propagated analytical uncertainty. *S. denticulatum* generally exhibited offsets below +0.05‰, but no clear species-specific fractionation pattern emerged from the complete dataset. Moreover, Δ^66^Zn_moss–solution_ showed no systematic relationship with the fraction of Zn adsorbed. The extent of Zn removal alone therefore does not explain the observed offsets and the differences in bonding of different isotopes likely stem from structural factors as examined in section 4.4 below.

Preferential retention of heavier Zn isotopes during adsorption from aqueous solution has been reported for a range of mineral and biological surfaces, although the magnitude of fractionation is generally greater than that observed here. Zn adsorption onto freshwater and marine diatoms enriches the cell-associated fraction in heavier isotopes, with fractionation of the order of several tenths of a per mil (Gélabert et al., 2006). Similarly, adsorption onto bacterial surfaces under high Zn-to-biomass conditions produced an average Δ^66^Zn_adsorbed–solution_ of approximately +0.46‰ (Kafantaris and Borrok, 2014), whereas short-term Zn adsorption onto phototrophic biofilms also resulted in preferential retention of heavier Zn isotopes by the biomass (Coutaud et al., 2014). In growing phototrophic biofilms, however, weak net Zn fractionation has been attributed to heavier-isotope adsorption combined with intracellular incorporation of lighter Zn (Coutaud et al., 2025). This biological pathway differs from the predominantly passive retention investigated with devitalized mosses.

Heavy-isotope enrichment has also been documented during Zn interaction with inorganic and organic sorbents. Sorption onto goethite and two-line ferrihydrite produced Δ^66^Zn_sorbed–aqueous_ values of approximately +0.29 and +0.53‰, respectively (Juillot et al., 2008), whereas adsorption onto amorphous Fe(III) oxyhydroxide yielded an enrichment of approximately +0.52‰ in the adsorbed phase (Balistrieri et al., 2008). In these mineral systems, preferential incorporation of heavier Zn isotopes has been related to differences in bond strength, Zn–O distance, and coordination environment between dissolved and surface-bound Zn. Recent isotope–XAS experiments further demonstrated contrasting isotope effects of adsorption and lattice substitution in Fe (hydr)oxides (Yan et al., 2023). Complexation with purified humic acid produced essentially no measurable fractionation below pH 6 but an enrichment of +0.24 ± 0.06‰ in organically bound Zn at higher pH, which was associated with increasing occupation of high-affinity binding sites (Jouvin et al., 2009). Together, these studies demonstrate that Zn isotope fractionation at environmental interfaces may depend on ligand identity, binding-site affinity, pH, aqueous speciation, and local coordination environment.

### 4.4. Testing whether bulk Zn coordination predicts isotope fractionation

A structural control on Zn isotope fractionation might therefore also be expected for mosses. Aqueous Zn^2+^ is predominantly hydrated in a six-coordinate environment, whereas adsorption and complexation at biological interfaces can produce lower-coordinate or more strongly bonded Zn complexes. Such changes have been invoked to explain preferential stabilization of heavier Zn isotopes at biological and mineral surfaces (Gélabert et al., 2006; Balistrieri et al., 2008; Juillot et al., 2008). However, the present results do not support a simple relationship between average first-shell structure and isotope fractionation. For samples with paired isotopic and spectroscopic measurements (*n* = 9), Δ^66^Zn_moss–solution_ was not significantly correlated with either the EXAFS-derived Zn–O distance (*R*; Spearman ρ = −0.04, *p* = 0.915) or coordination number (*N*; ρ = 0.14, *p* = 0.723). The Debye–Waller factor was likewise not significantly related to the isotopic offset. Thus, although local coordination changes may contribute to isotope discrimination at individual binding sites, the present data do not establish a predictive relationship between average *R* or *N* and bulk fractionation.

This lack of a significant correlation may reflect the heterogeneous surface sites and Zn-binding environments described in Sections 4.1 and 4.2. EXAFS-derived *R* and *N* represent averages across heterogeneous Zn populations rather than the structure of a single isotopically active complex. High-affinity sites unresolved by bulk XAS could contribute disproportionately to fractionation, while more abundant sites only weakly fractionate Zn isotopes. This interpretation is consistent with the contrasting isotope effects of organic-matter binding sites of differing affinities reported by Jouvin et al. (2009).

Limited compositional variability within the paired isotope–XAS subset further constrains identification of the binding environments that potentially control isotope fractionation. Cellulose-like environments dominated, whereas phosphate- and Zn–S-associated contributions were generally absent or below the practical resolution of linear-combination fitting. Relationships between Δ^66^Zn_moss–solution_ and cellulose-like or acetate-associated fractions were not significant by rank correlation. No single XANES-resolved component therefore emerged as a statistically supported predictor. Several factors may instead act jointly, including aqueous Zn speciation, ligand identity, adsorption mechanism, coordination chemistry, structural distortion and binding-site affinity (Duan et al., 2025).

The comparatively small moss isotope offsets may therefore integrate different site-specific isotope effects rather than reflect uniformly weak discrimination. In a mineral analogue, combined XAS, density functional theory and surface complexation modeling distinguished opposite Zn isotope effects at birnessite vacancy and edge sites (Wang et al., 2023). This illustrates how site contributions can combine, but does not imply lack of measurable isotopic offset in mosses. Resolving individual-site contributions requires experiments isolating changes in ligand distribution and Zn coordination across controlled pH and Zn-loading ranges.

### 4.5. Translating laboratory retention into isotope-based biomonitoring

The adsorption experiments constrain Zn retention by devitalized mosses in simplified aqueous systems rather than atmospheric accumulation involving particle interception, competing solutes and repeated wetting–drying. Rapid attainment of a concentration steady state does not, by itself, establish isotopic equilibrium. The small paired isotope–XAS dataset (*n* = 9), its limited compositional variability and the bulk-averaging nature of XAS restrict detection of individual binding-site controls. Non-significant correlations therefore do not exclude structural controls. Moreover, the comparison between native *S. denticulatum* and cultivated *S. palustre* combines species and cultivation differences, preventing isolation of their individual effects.

Nevertheless, the overlapping Zn isotope compositions reported for field-grown *Sphagnum* and *in vitro*-cultivated *S. palustre* clones are encouraging for biomonitoring, despite measurable differences in surface area, proton-reactive site density and Zn adsorption capacity. This similarity should not be interpreted as proof of equivalent moss–solution fractionation or the absence of cultivation-related bias. Together with their high adsorption capacity and standardized production, however, these observations support further evaluation of bioreactor-grown *Sphagnum* clones (Parsons et al., 2026) as alternatives to field-collected material. Controlled cultivation could improve comparability among monitoring sites and campaigns by providing material with a reproducible cultivation and pre-exposure history.

Recent field applications illustrate the potential of combining moss metal concentrations with isotope fingerprints. Jiao et al. (2026) used moss Zn isotopes to constrain industrial and combustion inputs across the Himalayas, assuming negligible fractionation during moss uptake. Aderinboye et al. (2026) demonstrated the complementary value of moss Pb isotope measurements for identifying aviation-fuel contributions near airports. However, this Pb application does not establish isotopic preservation for Zn. The small retention-related Zn offsets observed here provide a laboratory constraint on the assumption of isotopic preservation. Their significance for source discrimination should be evaluated relative to propagated analytical uncertainty and differences between source signatures, expressed on a common isotope-reference scale. The experimental offsets should not be applied as a universal correction to field measurements.

Field validation should therefore compare standardized native and cloned moss bags with co-located aerosol and wet-deposition samples, measuring Zn concentrations and isotope compositions before and after exposure. Initial biomass Zn, field blanks, soluble versus particle-associated inputs, and retention or loss during contrasting weather conditions should be considered. In particular, mixing of pre-existing and newly accumulated Zn must be distinguished from fractionation during retention. Such comparisons would establish whether cloned mosses preserve environmentally meaningful source contrasts and under which conditions retention modifies the recorded signal. Until then, the present results support their potential for isotope-based biomonitoring, but not quantitative preservation of atmospheric Zn isotope signatures under all exposure conditions.

## CONCLUSION

Zn retention by mosses was rapid, strongly pH-dependent, and predominantly surface-controlled. XANES and EXAFS revealed heterogeneous, mainly O-coordinated binding environments within the polysaccharide-rich cell-wall matrix, with variable phosphate and acetate contributions and minor Zn–S coordination in some samples. Moss-associated Zn was slightly enriched in heavy isotopes relative to solution (Δ⁶⁶Zn_moss–solution_ ranged from +0.01 to +0.09‰). Fractionation was not systematically related to the proportion of Zn adsorbed, Zn–O bond distance, or coordination number. These results do not support a simple structural control based on a single XAS-derived parameter and suggest that the isotopic signal may instead integrate the effects of multiple binding environments. Despite differences in surface properties and adsorption capacity, native and *in vitro*-cultivated *Sphagnum* showed no systematic difference in isotope fractionation within analytical resolution. Together with their high adsorption capacity and standardized production, these findings support bioreactor-grown clones as an alternative to field-collected moss for biomonitoring. However, atmospheric exposure experiments are needed before assuming that moss bags quantitatively preserve atmospheric Zn isotope signatures. Further paired isotope–XAS studies across controlled pH and Zn-loading ranges could help resolve the contributions of individual binding environments and improve interpretation of Zn accumulation in moss biomonitoring.

## ACKNOWLEDGMENTS

This work was supported by the MOSSCLONE project (FP7-ENV.2011.3.1.9-1, Grant Agreement No. 282952) of the European Union Seventh Framework Programme (FP7) for Research and Technological Development and by the Deutsche Forschungsgemeinschaft (DFG, German Research Foundation) under Germany’s Excellence Strategy (CIBSS – EXC-2189 – Project ID 39039984). The authors gratefully acknowledge the European Synchrotron Radiation Facility (ESRF, Grenoble, France) and the BM25-SpLine Spanish CRG beamline for providing synchrotron radiation facilities and assistance during the Zn K-edge XAS measurements.

## Declaration of generative AI and AI-assisted technologies in the manuscript preparation process

During preparation of the final version of the manuscript, the authors used OpenAI (Astra 6.0 Pro) to assist in improving the clarity and readability of author-prepared text and checking the manuscript for completeness and compliance with the journal’s guidelines. All AI-assisted outputs were critically reviewed, independently verified, and edited by the authors before inclusion in the manuscript. The authors take full responsibility for the accuracy, integrity, and content of the article.

## REFERENCES

Aceto, M., Abollino, O., Conca, R., Malandrino, M., Mentasti, E., Sarzanini, C., 2003. The use of mosses as environmental metal pollution indicators. Chemosphere 50, 333–342.

Aderinboye, O.E., Shiel, A.E., Stalford, L., Murphy, C.J., Atkinson, D.B., 2026. Moss biomonitoring and isotopic signatures reveal aviation fuel as local lead source near Oregon airports. Communications Sustainability 1, 128. DOI: 10.1038/s44458-026-00130-x.

Ares, A., Aboal, J.R., Carballeira, A., Giordano, S., Adamo, P., Fernández, J.A., 2012. Moss bag biomonitoring: A methodological review. Science of the Total Environment 432, 143–158.

Balistrieri, L.S., Borrok, D.M., Wanty, R.B., Ridley, W.I., 2008. Fractionation of Cu and Zn isotopes during adsorption onto amorphous Fe(III) oxyhydroxide: Experimental mixing of acid rock drainage and ambient river water. Geochimica et Cosmochimica Acta 72, 311– 328.

Beike, A.K., Spagnuolo, V., Lüth, V., Steinhart, F., Ramos-Gómez, J., Krebs, M., Adamo, P., Rey-Asensio, A.I., Fernández, J.A., Giordano, S., Decker, E.L., Reski, R., 2015. Clonal in vitro propagation of peat mosses (*Sphagnum* L.) as novel green resources for basic and applied research. Plant Cell, Tissue and Organ Culture 120, 1037–1049.

Capozzi, F., Giordano, S., Aboal, J.R., Adamo, P., Bargagli, R., Boquete, T., Di Palma, A., Real, C., Reski, R., Spagnuolo, V., Steinbauer, K., Tretiach, M., Varela, Z., Zechmeister, H., Fernandez, J.A., 2016. Best options for the exposure of traditional and innovative moss bags: A systematic evaluation in three European countries. Environmental Pollution 214, 362–373.

Capozzi, F., Adamo, P., Di Palma, A., Aboal, J.R., Bargagli, R., Fernández, J.A., López-Mahía, P., Reski, R., Tretiach, M., Spagnuolo, V., Giordano, S., 2017. *Sphagnum palustre* clone vs native *Pseudoscleropodium purum*: A first trial in the field to validate the future of the moss bag technique. Environmental Pollution 225, 323–328.

Cloquet, C., Carignan, J., Libourel, G., 2006. Isotopic composition of Zn and Pb atmospheric depositions in an urban/periurban area of northeastern France. Environmental Science & Technology 40, 6594–6600.

Cloquet, C., Carignan, J., Lehmann, M.F., Vanhaecke, F., 2008. Variation in the isotopic composition of zinc in the natural environment and the use of zinc isotopes in biogeosciences: A review. Analytical and Bioanalytical Chemistry 390, 451–463.

Clymo, R.S., 1963. Ion exchange in *Sphagnum* and its relation to bog ecology. Annals of Botany 27, 309–324.

Coutaud, A., Méheut, M., Viers, J., Rols, J.L., Pokrovsky, O.S., 2014. Zinc isotope fractionation during its interaction with phototrophic biofilm. Chemical Geology 390, 46–60.

Coutaud, M., Viers, J., Rols, J.-L., Pokrovsky, O.S., 2025. Copper and zinc isotope fractionation during phototrophic biofilm growth. Science of the Total Environment 960, 178371. DOI: 10.1016/j.scitotenv.2025.178371.

Di Palma, A., Crespo Pardo, D., Spagnuolo, V., Adamo, P., Bargagli, R., Cafasso, D., Capozzi, F., Aboal, J.R., González, A.G., Pokrovsky, O.S., Beike, A.K., Reski, R., Tretiach, M., Varela, Z., Giordano, S., 2016. Molecular and chemical characterization of a *Sphagnum palustre* clone: Key steps towards a standardized and sustainable moss bag technique. Ecological Indicators 71, 388–397.

Di Palma, A., Pallozzi, E., Gonzalez, A.G., Pokrovsky, O.S., Reski, R., Glime, J., Capozzi, F., Kavasi, N., Calfapietra, C., 2026. Linking biosurface reactivity and photosynthesis to investigate *Sphagnum* mosses as ecosystems engineers. New Phytologist, 252, 69–86, DOI: 10.1111/nph.71447.

Duan, Y., Wang, Z., Gou, W., Wang, Z., Li, Q., Li, W., 2025. Stable zinc isotopes as tracers in environmental geochemistry. Earth-Science Reviews 269, 105185.

Fernández, J.A., Boquete, M.T., Carballeira, A., Aboal, J.R., 2015. A critical review of protocols for moss biomonitoring of atmospheric deposition: Sampling and sample preparation. Science of the Total Environment 517, 132–150.

Fomina, M., Charnock, J.M., Hillier, S., Alexander, I.J., Gadd, G.M., 2006. Zinc phosphate transformations by the *Paxillus involutus*/pine ectomycorrhizal association. Microbial Ecology 52, 322–333.

Fomina, M., Charnock, J.M., Bowen, A.D., Gadd, G.M., 2007. X-ray absorption spectroscopy (XAS) of toxic metal mineral transformations by fungi. Environmental Microbiology 9, 308–321.

Gélabert, A., Pokrovsky, O.S., Viers, J., Schott, J., Boudou, A., Feurtet-Mazel, A., 2006. Interaction between zinc and freshwater and marine diatom species: Surface complexation and Zn isotope fractionation. Geochimica et Cosmochimica Acta 70, 839–857.

González, A.G., Pokrovsky, O.S., 2014. Metal adsorption on mosses: Toward a universal adsorption model. Journal of Colloid and Interface Science 415, 169–178.

González, A.G., Pokrovsky, O.S., Beike, A.K., Reski, R., Di Palma, A., Adamo, P., Giordano, S., Fernández, J.A., 2016. Metal and proton adsorption capacities of natural and cloned *Sphagnum* mosses. Journal of Colloid and Interface Science 461, 326–334.

Guiné, V., Spadini, L., Sarret, G., Muris, M., Delolme, C., Gaudet, J.-P., Martins, J.M.F., 2006. Zinc sorption to three Gram-negative bacteria: Combined titration, modeling, and EXAFS study. Environmental Science & Technology 40, 1806–1813.

Harmens, H., Norris, D.A., Steinnes, E., Kubin, E., Piispanen, J., Alber, R., Aleksiayenak, Y., Blum, O., Coşkun, M., Dam, M., De Temmerman, L., Fernández, J.A., Frolova, M., Frontasyeva, M., González-Miqueo, L., Grodzińska, K., Jeran, Z., Korzekwa, S., Krmar, M., Kvietkus, K., Leblond, S., Liiv, S., Magnússon, S.H., Mankovská, B., Pesch, R., Rühling, Å., Santamaria, J.M., Schröder, W., Spiric, Z., Suchara, I., Thöni, L., Urumov, V., Yurukova, L., Zechmeister, H.G., 2010. Mosses as biomonitors of atmospheric heavy metal deposition: Spatial patterns and temporal trends in Europe. Environmental Pollution 158, 3144–3156.

Heck, M.A., Lüth, V.M., van Gessel, N., Krebs, M., Kohl, M., Prager, A., Joosten, H., Decker, E.L., Reski, R., 2021a. Axenic in-vitro cultivation of 19 peat moss (*Sphagnum* L.) species as a resource for basic biology, biotechnology and paludiculture. New Phytologist 229, 861–876.

Heck, M.A., Melkova, I., Posten, C., Decker, E.L., Reski, R., 2021b. Medium optimization for biomass production of three peat moss (*Sphagnum* L.) species using fractional factorial design and response surface methodology. Bioresource Technology Reports 15, 100729.

Jiao, X., Dong, Z., Shao, Y., Baccolo, G., Li, F., Wu, R., 2026. Zinc isotopes reveal cross-border transport of coal combustion pollutants across the Himalayas during the summer monsoon. Geophysical Research Letters 53, e2026GL122536. DOI: 10.1029/2026GL122536.

Jouvin, D., Louvat, P., Juillot, F., Maréchal, C.N., Benedetti, M.F., 2009. Zinc isotopic fractionation: Why organic matters. Environmental Science & Technology 43, 5747–5754.

Juillot, F., Maréchal, C., Ponthieu, M., Cacaly, S., Morin, G., Benedetti, M.F., Hazemann, J.L., Proux, O., Guyot, F., 2008. Zn isotopic fractionation caused by sorption on goethite and 2-lines ferrihydrite. Geochimica et Cosmochimica Acta 72, 4886–4900.

Kafantaris, F.-C.A., Borrok, D.M., 2014. Zinc isotope fractionation during surface adsorption and intracellular incorporation by bacteria. Chemical Geology 366, 42–51.

Kłos, A., Gordzielik, E., Jóźwiak, M.A., Rajfur, M., 2014. Sorption of cadmium and zinc in selected species of epigeic mosses. Bulletin of Environmental Contamination and Toxicology 92, 323–328.

Maréchal, C.N., Télouk, P., Albarède, F., 1999. Precise analysis of copper and zinc isotopic compositions by plasma-source mass spectrometry. Chemical Geology 156, 251–273.

Parsons, J., Decker, E.L., Reski, R., 2026. Multiplication of peat moss (*Sphagnum* L.) species for climate action. Journal of Experimental Botany 77, 4087–4097.

Pokrovsky, O.S., Pokrovski, G.S., Gélabert, A., Schott, J., Boudou, A., 2005. Speciation of Zn associated with diatoms using X-ray absorption spectroscopy. Environmental Science & Technology 39, 4490–4498.

Popper, Z.A., Fry, S.C., 2003. Primary cell wall composition of bryophytes and charophytes. Annals of Botany 91, 1–12.

Reski, R., 1998. Development, genetics and molecular biology of mosses. Botanica Acta 111, 1–15.

Rühling, Å., Tyler, G., 1971. Regional differences in the deposition of heavy metals over Scandinavia. Journal of Applied Ecology 8, 497–507.

Viers, J., Prokushkin, A.S., Pokrovsky, O.S., Auda, Y., Kirdyanov, A.V., Beaulieu, E., Zouiten, C., Oliva, P., Dupré, B., 2013. Seasonal and spatial variability of elemental concentrations in boreal forest larch foliage of Central Siberia on continuous permafrost. Biogeochemistry 113, 435–449.

Viers, J., Prokushkin, A.S., Pokrovsky, O.S., Kirdyanov, A.V., Zouiten, C., Chmeleff, J., Chabaux, F., Oliva, P., Dupré, B., 2015. Zn isotope fractionation in a pristine larch forest on permafrost-dominated soils in Central Siberia. Geochemical Transactions, 16(1), 3, doi: 10.1186/s12932-015-0018-0.

Wang, Z., Peacock, C., Kwon, K.D., Gu, X., Feng, X., Li, W., 2023. Site-specific isotope fractionation during Zn adsorption onto birnessite: Insights from X-ray absorption spectroscopy, density functional theory and surface complexation modeling. Geochimica et Cosmochimica Acta 348, 68–84. DOI: 10.1016/j.gca.2023.03.006.

Wei, T., Dong, Z., Li, F., Kang, S., Qin, X., 2025. Quantifying the distribution and origins of aerosol zinc across the Northern Hemisphere using stable zinc isotopes: A review. Journal of Hazardous Materials 491, 137828.

