## Supplemental Information for "Zinc isotope fractionation in native and cloned mosses: implications for standardized pollution biomonitoring"

### Structural control on Zn isotope fractionation in native and cloned mosses: application for bioindicators

[3]Universidad Complutense de Madrid e IMDEA-Nanociencia

[4]Plant Biotechnology, Faculty of Biology, University of Freiburg, 79104 Freiburg, Germany

[5]CIBSS – Centre for Integrative Biological Signalling Studies, University of Freiburg, 79104 Freiburg, Germany

### SUPPLEMENTARY TEXT 1: DETAILED METHODS

#### Text S1.1. Chemicals and contamination control

All reagents were of analytical grade. Zinc nitrate [$Zn(NO_3)_2 \cdot xH_2O$; Sigma–Aldrich] was used as the $Zn^{2+}$ source. Solutions were prepared with ultrapure water (18.2 MΩ·cm; Milli-Q, Millipore), and 0.01 M $NaNO_3$ served as an inert background electrolyte to maintain constant ionic strength. For pH control, 2-(N-morpholino)ethanesulfonic acid (MES; Merck) was used at pH 5.5 and 4-(2-hydroxyethyl)-1-piperazineethanesulfonic acid (HEPES; Sigma–Aldrich) at pH 6.5. Solution pH was adjusted with small aliquots of standardized NaOH or $HNO_3$ (0.01–0.1 M; Merck Suprapur). Glassware and plasticware were acid-washed with 10% $HNO_3$ and thoroughly rinsed with ultrapure water before use to minimize trace-metal contamination.

#### Text S1.2. Moss pretreatment and adsorption procedures

Before adsorption experiments, mosses were rinsed three times with ultrapure water to remove attached particles and soluble salts, then devitalized at 120 °C overnight. Intact tissues were used without grinding or mechanical disaggregation to reproduce conditions relevant to moss-bag applications. Suspensions contained 1.0 g dry weight $L^{-1}$ and were continuously stirred with Teflon-coated magnetic stir bars. Gentle $N_2$ purging minimized dissolution of atmospheric $CO_2$.

In pH-edge experiments, pH was varied incrementally by adding standardized $HNO_3$ or NaOH (0.01–0.1 M), with at least 10 min of equilibration after each adjustment. In isotherm experiments, suspensions were equilibrated for 10 min after each Zn addition, based on preliminary kinetic experiments demonstrating rapid attainment of adsorption equilibrium. Following equilibration, suspensions were filtered through single-use 0.45-µm cellulose acetate membranes (Sartorius). Filtrates were acidified with ultrapure $HNO_3$ before dissolved-Zn analysis. Procedural blanks and moss-leachate controls without added Zn were processed in parallel.

#### Text S1.3. Moss provenance and surface characterization

The four field-collected moss taxa originated from non-urban sites in Galicia, NW Spain, away from direct pollution sources. The *S. palustre* clones 2a and 12a were cultivated *in vitro* at the University of Freiburg, Germany, within the EU-FP7 MossClone project. The clonal material provided a standardized alternative to field-collected *Sphagnum*, with a controlled cultivation history and reproducible physicochemical properties.

Surface acid–base properties were characterized by potentiometric titration. Proton adsorption was evaluated over a broad pH range, and the pH of zero net proton adsorption ($pH_{PZC}$) was derived from the titration curves. Proton-binding data were analyzed using the Linear Programming Model (LPM) to determine apparent acid–base equilibrium constants and the abundance of proton-reactive surface sites. Based on their apparent ranges, sites were tentatively assigned to carboxyl/phosphodiester, carboxyl, phosphoryl, amine and polyphenolic functional groups.

#### Text S1.4. Zn concentration and isotope analyses

*S1.4.1. Wet chemistry and analyses*

Dissolved Zn was determined using an Agilent 7500 ce ICP-MS. Samples were diluted with Milli-Q water to match the calibration range and acidified with bi-distilled $HNO_3$. Ten blanks were analyzed at the start of each run to calculate detection and quantification limits. Quality control included the certified reference material NRC-SLRS-6; the Zn quantification limit was 0.6 µg $L^{-1}$. Experimental solutions were also analyzed using an AAnalyst 400 AAS (Perkin Elmer) at 213.86 nm, with a quantification limit of 35 µg $L^{-1}$ and an uncertainty of 1.6%.

Moss samples were dried at 70 °C and digested in pre-cleaned Savillex vessels in a clean room at GET. Samples were resuspended in 1 mL of bi-distilled 15 N $HNO_3$, treated by ultrasonication for 30 min, and oxidized with 1 mL of $H_2O_2$ (30%; Merck superior quality). The remaining matrix was dissolved in 1.2 mL of 15 N $HNO_3$ plus 1.2 mL of HF (40%; Merck Suprapur), followed by re-oxidation with aqua regia. The treatment was conducted for 24 h at 70 °C, with evaporation at 70 °C between steps. Solution samples were evaporated at 70 °C.

Dried residues were resuspended in 1 mL of 7 N HCl + 0.001% $H_2O_2$. Zn was purified using 1.6 mL of AGMP-1 anion-exchange resin (Bio-Rad, USA). The resin was washed successively with ultrapure water and 0.5 N $HNO_3$ three times, then conditioned with 7 N HCl + 0.001% $H_2O_2$. Following sample loading and successive elution of the matrix, Cu and Fe, Zn was eluted with 0.5 N $HNO_3$. Zn recovery was checked by ICP-MS measurements before and after purification, and only samples yielding 95–100% recovery were retained for isotope analysis.

#### *S1.4.2. Zn isotope measurements and quality control*

##### Instrumentation and data acquisition

Zn isotope ratios were measured using a Neptune MC-ICP-MS (ThermoFinnigan, GET) and a P54 MC-ICP-MS (VG Elemental, ENS, Lyon, France). The Neptune sample-introduction system comprised a tandem quartz-glass spray chamber (cyclonic and standard Scott double-pass chambers) coupled to a low-flow PFA nebulizer. Argon flow rates were approximately 15, 1.2 and 0.6 L $min^{-1}$ for the coolant, nebulizer and auxiliary gas, respectively, and the radiofrequency power was 1300 W.

The Neptune multicollector module comprised eight movable Faraday cups and one fixed cup, with a maximum relative mass range of 17%, allowing simultaneous collection of $^{64}Zn$, $^{66}Zn$, $^{67}Zn$, $^{68}Zn$, $^{63}Cu$, $^{65}Cu$ and $^{62}Ni$ (Viers et al., 2015). After at least 3 h of warm-up and tuning, sequences were run using a CETAC ASX-100 autosampler. Between sample and standard measurements, rinsing used 0.05 N $HNO_3$ from two separate vials for 1 min each. Blanks were measured in one block of 10 cycles, and samples and standards in two blocks of 20 cycles, with 8 s per cycle.

##### Mass-bias correction and isotope reporting

Instrumental mass fractionation was corrected using sample–standard bracketing and Cu internal standardization (Viers et al., 2015). NIST 976 Cu was added to purified Zn fractions, and a mixture of NIST 976 Cu and JMC 3-0749 L Zn was used as the bracketing standard. Cu and Zn fractionation factors were checked during one analytical session and remained constant.

Zn isotope compositions were expressed relative to JMC batch 3-0749 L, a mono-elemental solution supplied by ENS, Lyon, and used by several laboratories as an isotopic standard rather than a certified reference material. Values were reported in delta notation (Viers et al., 2015):

$$\delta^{66}\mathrm{Zn} = \left[\frac{(^{66}\mathrm{Zn}/^{64}\mathrm{Zn})_s}{(^{66}\mathrm{Zn}/^{64}\mathrm{Zn})_{\mathrm{JMC}}} - 1\right] \times 1000\ (‰)$$

where *s* denotes the sample and JMC denotes the Zn standard solution.

##### Quality control and uncertainty

The $^{62}Ni$ signal was monitored to assess $^{64}Ni$ interference on $^{64}Zn$; no significant interference was detected. Zn isotope ratios were checked for consistency with theoretical mass-dependent fractionation. Clean-room blanks, an in-house biofilm standard and the lichen-powder reference material CRM-BCR 482 (standard no. 685, IRMM) were analyzed regularly. Internal precision for sample and standard measurements was 5–10 ppm (2σ std) for both $^{65}Cu/^{63}Cu$ and $^{66}Zn/^{64}Zn$.

Repeated measurements of the in-house biofilm standard yielded $\sigma/\sqrt{N}$ = 0.008‰ ($N$ = 7), where σ is the standard deviation. Each reported sample value represents the mean of 3–5 measurements, with uncertainty calculated as $\sigma/\sqrt{N}$ and a minimum assigned uncertainty of 0.008‰. Uncertainties in calculated isotopic separation factors were propagated from the measurement uncertainties. The dispersion of mean isotope offsets ($\Delta^{66}$Zn) and enrichment factors ($\varepsilon^{66}$Zn) was expressed as the standard deviation.

**Text S1.5. XAS sample handling, acquisition and fitting**

Zn K-edge measurements were performed at the BM25-SpLine Spanish CRG beamline of the European Synchrotron Radiation Facility (ESRF, Grenoble, France), using a Si(111) double-crystal monochromator. After laboratory adsorption, moss samples were briefly rinsed with ultrapure water to remove loosely associated Zn, blotted, immediately frozen in liquid nitrogen and stored at −80 °C until analysis. Samples were mounted in a cryostat and maintained at approximately 10 K to minimize thermal disorder and radiation-induced changes. XANES and EXAFS spectra were collected in fluorescence mode using a multi-element Ge solid-state detector. Energy was calibrated against metallic Zn foil, assigning its first K-edge inflection point to 9659 eV.

Reference compounds were measured under comparable conditions and included ZnO, ZnS, Zn phosphate, Zn acetate, Zn-associated cellulose, Zn citrate, $ZnCO_3$, $Zn_2SiO_4$ and aqueous $Zn^{2+}$ prepared from $Zn(NO_3)_2$ solutions. Spectra were processed in ATHENA by energy calibration, background subtraction, normalization and extraction of the EXAFS function. XANES spectra were analyzed by least-squares linear-combination fitting using the measured references to estimate the relative contributions of Zn coordination environments. EXAFS functions were Fourier transformed, and structural parameters were obtained in ARTEMIS by fitting FEFF-calculated scattering paths. First-shell models predominantly used Zn–O paths; Zn–S paths were added where sulfur coordination was independently indicated by XANES.

**Table S1.** Experimental conditions and Zn adsorption characteristics of native and *in vitro*-cultivated moss samples used for adsorption, isotope, and XAS analyses. Reference biomass of moss samples means intact natural biomass.

| Sample | Species | Type of study | $[Zn]_0$ mg/L | $[Zn]_{aq}$ mg/L | $[Zn]_{ads}$ mg/L | $\%[Zn]_{ads}$ | pH |
|---|---|---|---|---|---|---|---|
| **Native samples** | | | | | | | |
| Zn-1 | *Sphagnum sp.* | Long –term | 3.04 | 0.46 | 2.58 | 84.9 | 6.17 |
| Zn-2 | *Sphagnum sp.* | Long –term | 3.04 | 0.26 | 2.78 | 91.5 | 6.21 |
| Zn-3 | *Sphagnum sp.* | Long –term | 3.05 | 0.22 | 2.83 | 92.8 | 6.07 |
| Zn-4 | *Sphagnum sp.* | Long –term | 4.13 | 0.54 | 3.59 | 86.9 | 5.36 |
| Zn-5 | *Sphagnum sp.* | Long –term | 50.4 | 27.7 | 22.7 | 45.0 | 6.15 |
| Zn-6 | *Sphagnum sp.* | Long –term | 50.8 | 26.3 | 24.5 | 48.2 | 6.17 |
| Zn-7 | *Sphagnum sp.* | Long –term | 50.5 | 26.5 | 24.0 | 47.5 | 5.86 |
| Zn-8 | *Sphagnum sp.* | Long –term | 54.1 | 27.5 | 26.7 | 49.3 | 5.27 |
| Zn-9 | *Sphagnum sp.* | Long –term | 101 | 58.9 | 42.2 | 41.8 | 6.19 |
| Zn-10 | *Sphagnum sp.* | Long –term | 86.0 | 52.1 | 33.9 | 39.4 | 6.14 |
| Zn-11 | *Sphagnum sp.* | Long –term | 101 | 58.8 | 42.4 | 41.9 | 5.81 |
| Zn-12 | *Sphagnum sp.* | Long –term | 101 | 57.9 | 43.1 | 42.7 | 4.55 |
| Zn-13 | *Sphagnum sp.* | Long –term | 152 | 91.5 | 60.3 | 39.7 | 6.24 |
| Zn-14 | *Sphagnum sp.* | Long –term | 152 | 87.9 | 64.2 | 42.2 | 6.07 |
| Zn-15 | *Sphagnum sp.* | Long –term | 152 | 91.0 | 60.6 | 40.0 | 5.69 |
| Zn-16 | *Sphagnum sp.* | Long –term | 152 | 90.8 | 60.9 | 40.2 | 4.59 |
| Zn-17 | *Sphagnum sp.* | Long –term | 202 | 124 | 78.2 | 38.7 | 6.27 |
| Zn-18 | *Sphagnum sp.* | Long –term | 202 | 121 | 81.6 | 40.3 | 6.17 |
| Zn-19 | *Sphagnum sp.* | Long –term | 202 | 127 | 75.5 | 37.3 | 6.01 |
| Zn-20 | *Sphagnum sp.* | Long –term | 203 | 123 | 80.0 | 39.5 | 4.49 |
| Zn-21 | *Hypnum sp.* | Langmuirian | 232 | 154 | 78.4 | 33.7 | 6.76 |
| Zn-22 | *Sphagnum sp.* | Langmuirian | 237 | 152 | 84.4 | 35.7 | 6.79 |
| Zn-23 | *P. purum* | Langmuirian | 234 | 157 | 77.6 | 33.2 | 6.77 |
| Zn-24 | *B. rutabulum* | Langmuirian | 229 | 146 | 82.9 | 36.2 | 6.78 |
| Zn-25 | *Hypnum sp.* | pH-edge | 3.3 | 3.28 | 0.02 | 0.61 | 2.33 |
| Zn-26 | *Sphagnum sp.* | pH-edge | 3.3 | 3.30 | 0 | 0 | 2.04 |
| Zn-27 | *P. purum* | pH-edge | 3.3 | 3.25 | 0.05 | 1.52 | 2.13 |
| Zn-28 | *B. rutabulum* | pH-edge | 3.3 | 3.21 | 0.09 | 2.70 | 2.12 |
| Zn-29 | *Hypnum sp.* | pH-edge | 3.3 | 2.23 | 1.07 | 32.5 | 4.81 |
| Zn-30 | *Sphagnum sp.* | pH-edge | 3.3 | 1.91 | 1.40 | 42.3 | 4.86 |
| Zn-31 | *P. purum* | pH-edge | 3.3 | 1.90 | 1.40 | 42.4 | 4.76 |
| Zn-32 | *B. rutabulum* | pH-edge | 3.3 | 1.53 | 1.77 | 53.7 | 5.61 |
| Zn-33 | *Hypnum sp.* | pH-edge | 3.3 | 1.34 | 1.96 | 59.3 | 7.16 |
| Zn-34 | *Sphagnum sp.* | pH-edge | 3.3 | 1.70 | 1.60 | 48.6 | 7.20 |
| Zn-35 | *P. purum* | pH-edge | 3.3 | 1.51 | 1.80 | 54.4 | 7.14 |
| Zn-36 | *B. rutabulum* | pH-edge | 3.3 | 1.25 | 2.05 | 62.4 | 7.17 |
| Zn-37 | *Hypnum sp.* | pH-edge | 200 | 109 | 91.0 | 45.5 | 1.88 |
| Zn-38 | *Sphagnum sp.* | pH-edge | 200 | 116 | 84.3 | 42.2 | 2.25 |
| Zn-39 | *P. purum* | pH-edge | 200 | 119 | 80.8 | 40.4 | 2.23 |
| Zn-40 | *B. rutabulum* | pH-edge | 200 | 95.4 | 105 | 52.3 | 2.10 |

| | | | | | | | |
|---|---|---|---|---|---|---|---|
| Zn-47 | *Sphagnum sp.* | growth | 0.43 | 1.23 | -0.80* | -186* | |
| Zn-48 | *Sphagnum sp.* | growth | 3.73 | 4.01 | -0.28* | -7.5* | |
| Zn-49 | *Sphagnum sp.* | growth | 50.4 | 50.6 | -0.19* | -0.37* | |
| Zn-50 | *Sphagnum sp.* | growth | 150 | 154 | -3.09* | -2.1* | |
| Zn-51 | *Sphagnum sp.* | growth | 250 | 248 | 2.42 | 0.97* | |
| ***in vitro*-cultivated moss samples** | | | | | | | |
| Zn-41 | *Sphagnum sp.Clone C-2a* | Isotopes | 3.3 | 1.38 | 1.92 | 41.8 | 5.19 |
| Zn-42 | *Sphagnum sp. Clone C-2a* | Isotopes | 3.3 | 0.35 | 2.95 | 10.6 | 3.59 |
| Zn-43 | *Sphagnum sp. Clone C-2a* | Isotopes | 3.3 | 2.02 | 1.23 | 61.3 | 6.41 |
| Zn-44 | *Sphagnum sp. Clone C-12a* | Isotopes | 3.3 | 1.72 | 1.58 | 52.1 | 4.03 |
| Zn-45 | *Sphagnum sp. Clone C-12a* | Isotopes | 3.3 | 2.06 | 1.24 | 62.4 | 5.05 |
| Zn-46 | *Sphagnum sp. Clone C-12a* | Isotopes | 3.3 | 0.62 | 2.68 | 18.8 | 2.35 |
| **Reference samples** | | | | | | | |
| Zn-52 | *Hypnum sp.* | Intact biomass | --- | --- | --- | --- | --- |
| Zn-53 | *Sphagnum sp.* | Intact biomass | --- | --- | --- | --- | --- |
| Zn-54 | *P. purum* | Intact biomass | --- | --- | --- | --- | --- |
| Zn-55 | *B. rutabulum* | Intact biomass | --- | --- | --- | --- | --- |
| Zn-55 | *Cellulose* | pH-edge | 3.3 | 3.601 | -0.30 | -9.12 | 2.18 |
| Zn-55 | *Cellulose* | pH-edge | 3.3 | 0.065 | 3.24 | 98. | 10.9 |
| Zn-55 | *Cellulose* | pH-edge | 3.3 | 3.237 | 0.06 | 1.91 | 5.02 |

*Negative adsorption means Zn release from biomass

### Text to Table S1 — Surface properties and detailed Zn adsorption results

For the four field-collected mosses, the $pH_{PZC}$ was 5.01 ± 0.13 for *Hypnum* sp., 4.64 ± 0.10 for *S. denticulatum*, 4.96 ± 0.14 for *P. purum*, and 6.23 ± 0.25 for *B. rutabulum*. Among these species, *S. denticulatum* exhibited the lowest $pH_{PZC}$ and the highest density of proton-reactive surface sites (0.65 mmol $g^{-1}$), compared with 0.55, 0.49, and 0.48 mmol $g^{-1}$ for *P. purum*, *Hypnum* sp., and *B. rutabulum*, respectively (González and Pokrovsky, 2014). LPM analysis resolved five main classes of proton-reactive sites, tentatively assigned to phosphodiester, carboxyl, phosphoryl, amine, and polyphenolic functional groups according to their apparent values.

At an initial $Zn^{2+}$ concentration of 3.3 mg $L^{-1}$, adsorption was negligible under strongly acidic conditions (0–2.7% at pH 2.04–2.33), but increased markedly with increasing pH. At intermediate pH (4.76–5.61), 32.5–53.7% of the added $Zn^{2+}$ was adsorbed, whereas at pH 7.14–7.20 adsorption ranged from 48.6 to 62.2% (Table S1). Despite species-specific differences in the position and shape of the adsorption edges, all moss species displayed the same overall sigmoidal dependence of Zn retention on pH, consistent with the adsorption behavior previously reported for these materials (González and Pokrovsky, 2014).

In the adsorption-isotherm experiments conducted at pH 6.76–6.79 and initial $Zn^{2+}$ concentrations of approximately 229–237 mg $L^{-1}$, the fraction of $Zn^{2+}$ adsorbed ranged only from 33.2 to 36.2% among the four species (Table S1). Similarly, long-term experiments with *Sphagnum* showed a strong dependence of fractional $Zn^{2+}$ retention on metal loading: 84.9–92.8% of $Zn^{2+}$ was retained at initial concentrations of approximately 3–4 mg $L^{-1}$, whereas the adsorbed fraction decreased to approximately 37.3–49.3% at initial $Zn^{2+}$ concentrations of 50–203 mg $L^{-1}$.

The in vitro-cultivated *S. palustre* clones exhibited acid–base characteristics broadly similar to those of field-grown *Sphagnum* but differed quantitatively in their surface properties. Previous characterization showed mass-normalized binding-site densities of 1.56 and 1.31 mmol $g^{-1}$ for clones 2a and 12a, respectively, i.e., approximately 1.5–2 times higher than that of native *S. denticulatum* (González et al., 2016). The clones also exhibited substantially larger specific surface areas (24 ± 1 and 28 ± 1 $m^2$ $g^{-1}$ for clones 2a and 12a, respectively) than the natural material (10.8 ± 0.3 $m^2$ $g^{-1}$), together with an approximately two-fold higher $Zn^{2+}$ adsorption capacity.

**Table S2.** Zn concentrations, adsorption parameters, and stable Zn isotope compositions of moss biomass and corresponding aqueous solutions. Reference biomass of moss samples mean intact nature biomass.

| **Sample** | **Species** | **Type of sample** | **$[M]_{aq}$ mg/L** | **$[M]_{ad}$ mg/L** | **% $M_{ad}$** | **pH** | **Zn (mg/kg)** | **$\delta^{66}$Zn (‰)** | **Std** | **$\delta^{67}$Zn (‰)** | **Std** | **$\delta^{68}$Zn (‰)** | **Std** | **$\Delta^{66}$Zn, ‰ solid-liquid** |
|---|---|---|---|---|---|---|---|---|---|---|---|---|---|---|
| **Reference** | | | | | | | | | | | | | | |
| Iso-H1 | *Hypnum sp.* | Solid | --- | --- | --- | --- | 27.4 | 0.111 | 0.014 | 0.257 | 0.006 | 0.263 | 0.008 | |
| Iso-S1 | *Sphagnum sp.* | Solid | --- | --- | --- | --- | 36.0 | 0.442 | 0.002 | 0.731 | 0.018 | 0.922 | 0.009 | |
| Iso-P1 | *P. purum* | Solid | --- | --- | --- | --- | 39.5 | 0.077 | 0.018 | 0.191 | 0.037 | 0.169 | 0.040 | |
| Iso-B1 | *B. rutabulum* | Solid | --- | --- | --- | --- | 36.9 | 0.273 | 0.023 | 0.504 | 0.038 | 0.556 | 0.054 | |
| **Native Samples** | | | | | | | | | | | | | | |
| Iso-H2 | *Hypnum sp.* | Solid | 1.704 | 1.174 | 40.8 | 4.97 | 1450 | 0.136 | 0.005 | 0.282 | 0.011 | 0.310 | 0.008 | 0.039 |
| Iso-S2 | *Sphagnum sp.* | Solid | 1.554 | 1.317 | 45.9 | 4.72 | 1429 | 0.162 | 0.015 | | | | | 0.050 |
| Iso-P2 | *P. purum* | Solid | 1.681 | 1.196 | 41.6 | 5.16 | 1627 | 0.127 | 0.006 | 0.260 | 0.001 | 0.295 | 0.005 | 0.026 |
| Iso-B2 | *B. rutabulum* | Solid | 1.430 | 1.443 | 50.2 | 5.25 | 1948 | 0.115 | 0.028 | 0.302 | 0.027 | 0.262 | 0.030 | 0.013 |
| Iso-H3 | *Hypnum sp.* | Solid | 2.185 | 0.623 | 22.2 | 3.94 | 937000 | 0.129 | 0.003 | 0.271 | 0.00003 | 0.285 | 0.003 | 0.018 |
| Iso-S3 | *Sphagnum sp.* | Solid | 2.356 | 0.470 | 16.2 | 4.14 | 1015000 | 0.139 | 0.019 | 0.161 | 0.020 | 0.283 | 0.009 | 0.048 |
| Iso-P3 | *P. purum* | Solid | 2.256 | 0.571 | 20.2 | 4.16 | 914000 | 0.132 | 0.018 | 0.320 | 0.017 | 0.291 | 0.025 | 0.035 |
| Iso-B3 | *B. rutabulum* | Solid | 2.120 | 0.682 | 24.4 | 3.88 | 1209 | 0.135 | 0.008 | 0.333 | 0.002 | 0.313 | 0.003 | 0.035 |
| Iso-H4 | *Hypnum sp.* | Solid | 0.807 | 1.51 | 65.1 | 7.10 | | 0.168 | 0.045 | | | | | 0.094 |
| Iso-S4 | *Sphagnum sp.* | Solid | 0.708 | 1.611 | 69.5 | 7.09 | 2320000 | 0.140 | 0.053 | 0.157 | 0.076 | 0.206 | 0.057 | 0.016 |
| Iso-P4 | *P. purum* | Solid | 0.861 | 1.34 | 60.8 | 7.30 | 2408000 | 0.083 | 0.032 | 0.221 | 0.047 | 0.255 | 0.023 | 0.004 |
| Iso-B4 | *B. rutabulum* | Solid | 0.871 | 1.47 | 62.8 | 7.05 | 2492000 | 0.092 | 0.056 | 0.243 | 0.101 | 0.286 | 0.027 | 0.016 |
| XAS-1 | *Hypnum sp.* | Solid | 3.28 | 0.02 | 0.6 | 2.33 | | 0.154 | 0.036 | 0.160 | 0.044 | 0.251 | 0.060 | 0.079 |
| XAS-2 | *Sphagnum sp.* | Solid | 3.29 | 0.01 | 0.3 | 2.04 | | 0.145 | 0.020 | 0.264 | 0.035 | 0.326 | 0.040 | 0.046 |
| XAS-3 | *P. purum* | Solid | 3.25 | 0.05 | 1.5 | 2.13 | | 0.113 | 0.052 | 0.278 | 0.072 | 0.258 | 0.099 | 0.020 |
| XAS-4 | *B. rutabulum* | Solid | 3.21 | 0.09 | 2.7 | 2.12 | | 0.147 | 0.010 | 0.275 | 0.003 | 0.293 | 0.033 | 0.051 |
| XAS-5 | *Hypnum sp.* | Solid | 2.23 | 1.07 | 32.4 | 4.81 | | 0.093 | 0.0295 | 0.218 | 0.043 | 0.193 | 0.060 | 0.029 |
| XAS-6 | *Sphagnum sp.* | Solid | 1.91 | 1.39 | 42.1 | 4.86 | | 0.106 | 0.038 | 0.198 | 0.042 | 0.242 | 0.077 | 0.037 |
| XAS-7 | *P. purum* | Solid | 1.9 | 1.4 | 42.4 | 4.76 | | 0.091 | 0.017 | 0.184 | 0.022 | 0.233 | 0.028 | -0.029 |
| XAS-8 | *B. rutabulum* | Solid | 1.53 | 1.77 | 53.6 | 5.61 | | 0.127 | 0.0028 | 0.194 | 0.001 | 0.269 | 0.002 | 0.059 |
| XAS-16 | *Hypnum sp.* | Solid | 1.34 | 1.96 | 59.4 | 7.16 | | 0.102 | 0.023 | 0.160 | 0.040 | 0.187 | 0.040 | 0.057 |

| | | | | | | | | | | | | | | |
|---|---|---|---|---|---|---|---|---|---|---|---|---|---|---|
| XAS-17 | *Sphagnum sp.* | Solid | 1.7 | 1.6 | 48.5 | 7.2 | | 0.138 | 0.289 | 0.440 | 0.396 | 0.258 | | 0.033 |
| Iso-H2 | *Hypnum sp.* | Liquid | 1.704 | 1.17 | 40.8 | 4.97 | | 0.097 | 0.011 | 0.144 | 0.022 | 0.197 | 0.032 | |
| Iso-S2 | *Sphagnum sp.* | Liquid | 1.554 | 1.32 | 45.9 | 4.72 | | 0.112 | 0.007 | 0.191 | 0.020 | 0.228 | 0.022 | |
| Iso-P2 | *P. purum* | Liquid | 1.681 | 1.196 | 41.6 | 5.16 | | 0.101 | 0.004 | 0.166 | 0.008 | 0.202 | 0.011 | |
| Iso-B2 | *B. rutabulum* | Liquid | 1.430 | 1.443 | 50.2 | 5.25 | | 0.086 | 0.003 | 0.162 | 0.003 | 0.200 | 0.002 | |
| Iso-H3 | *Hypnum sp.* | Liquid | 2.185 | 0.623 | 22.2 | 3.94 | | 0.111 | 0.002 | 0.188 | 0.003 | 0.228 | 0.005 | |
| Iso-S3 | *Sphagnum sp.* | Liquid | 2.356 | 0.470 | 16.6 | 4.14 | | 0.091 | 0.002 | 0.156 | 0.002 | 0.204 | 0.003 | |
| Iso-P3 | *P. purum* | Liquid | 2.256 | 0.571 | 20.2 | 4.16 | | 0.097 | 0.001 | 0.178 | 0.002 | 0.195 | 0.001 | |
| Iso-B3 | *B. rutabulum* | Liquid | 2.120 | 0.682 | 24.4 | 3.88 | | 0.100 | 0.019 | 0.149 | 0.037 | 0.205 | 0.041 | |
| Iso-H4 | *Hypnum sp.* | Liquid | 0.807 | 1.506 | 65.1 | 7.10 | | 0.074 | 0.012 | 0.115 | 0.012 | 0.134 | 0.011 | |
| Iso-S4 | *Sphagnum sp.* | Liquid | 0.708 | 1.611 | 69.5 | 7.09 | | 0.124 | 0.007 | 0.217 | 0.008 | 0.259 | 0.014 | |
| Iso-P4 | *P. purum* | Liquid | 0.861 | 1.34 | 60.8 | 7.30 | | 0.079 | 0.002 | 0.140 | 0.003 | 0.169 | 0.012 | |
| Iso-B4 | *B. rutabulum* | Liquid | 0.871 | 1.47 | 62.8 | 7.05 | | 0.076 | 0.007 | 0.141 | 0.006 | 0.169 | 0.013 | |
| Iso-H5 | *Hypnum sp.* | Liquid | 0.03 | 0.014 | 46.7 | 6.43 | | -0.010 | 0.012 | 0.090 | 0.008 | 0.054 | 0.039 | |
| Iso-S5 | *Sphagnum sp.* | Liquid | 0.03 | 0.014 | 46.3 | 5.31 | | -0.048 | 0.007 | 0.484 | 0.034 | 0.053 | 0.030 | |
| Iso-P5 | *P. purum* | Liquid | 0.03 | 0.015 | 50.6 | 4.98 | | 0.036 | 0.042 | 0.130 | 0.072 | 0.069 | 0.121 | |
| Iso-B5 | *B. rutabulum* | Liquid | 0.03 | 0.014 | 46.9 | 4.77 | | 0.036 | 0.067 | 0.165 | 0.090 | 0.027 | 0.141 | |
| Iso-H6 | *Hypnum sp.* | Liquid | 0.2 | 0.108 | 53.8 | 5.97 | | 0.097 | 0.085 | 0.124 | 0.132 | 0.135 | 0.159 | |
| Iso-S6 | *Sphagnum sp.* | Liquid | 0.2 | 0.102875 | 51.4 | 5.15 | | 0.058 | 0.005 | 0.052 | 0.019 | 0.174 | 0.017 | |
| Iso-P6 | *P. purum* | Liquid | 0.2 | 0.120 | 60.1 | 5.09 | | 0.048 | 0.023 | 0.155 | 0.056 | 0.231 | 0.054 | |
| Iso-B6 | *B. rutabulum* | Liquid | 0.2 | 0.105 | 52.7 | 5.24 | | 0.079 | 0.016 | 0.117 | 0.024 | 0.213 | 0.020 | |
| Iso-H7 | *Hypnum sp.* | Liquid | 1.0 | 0.613 | 61.3 | 4.92 | | 0.093 | 0.023 | 0.143 | 0.034 | 0.231 | 0.042 | |
| Iso-S7 | *Sphagnum sp.* | Liquid | 1.0 | 0.697 | 69.7 | 4.7 | | 0.031 | 0.006 | 0.600 | 0.081 | 0.206 | 0.013 | |
| Iso-P7 | *P. purum* | Liquid | 1.0 | 0.653 | 65.3 | 4.72 | | 0.065 | 0.001 | 0.112 | 0.082 | 0.205 | 0.089 | |
| Iso-B7 | *B. rutabulum* | Liquid | 1.0 | 0.540 | 54.1 | 5.39 | | 0.013 | 0.053 | 0.153 | 0.089 | 0.147 | 0.102 | |
| XAS-1 | *Hypnum sp.* | Liquid | 3.28 | 0.02 | 0.6 | 2.33 | | 0.075 | 0.043 | 0.126 | 0.020 | 0.145 | 0.077 | |

| | | | | | | | | | | | | | |
|---|---|---|---|---|---|---|---|---|---|---|---|---|---|
| XAS-2 | *Sphagnum sp.* | Liquid | 3.29 | 0.01 | 0.3 | 2.04 | | 0.0985 | 0.019 | 0.175 | 0.034 | 0.221 | 0.046 |
| XAS-3 | *P. purum* | Liquid | 3.25 | 0.05 | 1.5 | 2.13 | | 0.094 | 0.023 | 0.185 | 0.041 | 0.212 | 0.059 |
| XAS-4 | *B. rutabulum* | Liquid | 3.21 | 0.09 | 2.7 | 2.12 | | 0.095 | | 0.173 | | 0.197 | |
| XAS-5 | *Hypnum sp.* | Liquid | 2.23 | 1.07 | 32.4 | 4.81 | | 0.064 | 0.070 | 0.123 | | 0.100 | 0.121 |
| XAS-6 | *Sphagnum sp.* | Liquid | 1.91 | 1.39 | 42.1 | 4.86 | | 0.068 | 0.059 | 0.064 | 0.082 | 0.131 | 0.094 |
| XAS-7 | *P. purum* | Liquid | 1.9 | 1.4 | 42.4 | 4.76 | | 0.120 | 0.008 | 0.158 | 0.007 | 0.200 | 0.002 |
| XAS-8 | *B. rutabulum* | Liquid | 1.53 | 1.77 | 53.6 | 5.61 | | 0.068 | 0.032 | 0.154 | 0.073 | 0.160 | 0.073 |
| XAS-16 | *Hypnum sp.* | Liquid | 1.34 | 1.96 | 59.4 | 7.16 | | 0.044 | 0.011 | 0.054 | 0.016 | 0.090 | 0.013 |
| XAS-17 | *Sphagnum sp.* | Liquid | 1.7 | 1.6 | 48.5 | 7.2 | | 0.105 | 0.018 | 0.212 | 0.030 | 0.215 | 0.030 |
| XAS-18 | *P. purum* | Liquid | 1.51 | 1.79 | 54.2 | 7.14 | | 0.159 | 0.044 | 0.281 | 0.066 | 0.323 | 0.081 |
| XAS-19 | *B. rutabulum* | Liquid | 1.25 | 2.05 | 62.1 | 7.17 | | 0.130 | 0.005 | 0.244 | 0.027 | 0.258 | 0.009 |
| ***in vitro*-cultivated moss samples** | | | | | | | | | | | | | |
| C2a-1 | *Sphagnum sp.* | Liquid | 1.4 | 1.92 | 41.8 | 5.19 | | 0.106 | 0.002 | 0.208 | 0.004 | 0.204 | 0.010 |
| C2a-2 | *Sphagnum sp.* | Liquid | 0.4 | 2.95 | 10.6 | 3.59 | | 0.108 | 0.061 | 0.149 | 0.075 | 0.186 | 0.121 |
| C2a-3 | *Sphagnum sp.* | Liquid | 2.0 | 1.23 | 61.3 | 6.41 | | 0.072 | 0.047 | 0.141 | 0.056 | 0.145 | 0.081 |
| C12a-1 | *Sphagnum sp.* | Liquid | 1.7 | 1.58 | 52.1 | 4.03 | | 0.110 | 0.020 | 0.195 | 0.028 | 0.227 | 0.051 |
| C12a-2 | *Sphagnum sp.* | Liquid | 2.1 | 1.24 | 62.4 | 5.05 | | 0.091 | 0.047 | 0.261 | 0.056 | 0.246 | 0.094 |
| C12a-3 | *Sphagnum sp.* | Liquid | 0.6 | 2.68 | 18.8 | 2.35 | | 0.102 | 0.061 | 0.186 | 0.114 | 0.216 | 0.140 |
| **Standard Samples** | | | | | | | | | | | | | |
| Standard | Zn-nitrate | Liquid | | | | | | 0.135 | 0.014 | 0.166 | 0.011 | 0.242 | 0.021 |
| Standard | Zn-nitrate | Liquid | | | | | | 0.095 | 0.049 | 0.148 | 0.099 | 0.173 | 0.139 |

**Text to Table S2 — Isotope notation and detailed Zn isotope comparisons**

The recommended delta notation and the solid–solution isotopic offset were defined as follows:

$$\delta^{66}Zn = [((^{66}Zn/^{64}Zn)_{sample} / (^{66}Zn/^{64}Zn)_{STD}) - 1] \times 1000\ (‰) \quad (S1)$$

$$\Delta^{66}Zn_{solid\text{-}solution} = \delta^{66}Zn_{solid} - \delta^{66}Zn_{solution} \quad (S2)$$

The enrichment of the moss-associated Zn in the heavy isotope was observed over a broad range of Zn adsorption, from less than 5% to approximately 70% of the initially available Zn (Fig. 2). For example, values of 0.039, 0.026, and 0.013‰ were obtained for *Hypnum* sp., *P. purum*, and *B. rutabulum*, respectively, under conditions where approximately 41–50% of the $Zn^{2+}$ was adsorbed. At lower adsorption levels (approximately 17–24%), isotopic offsets ranged from 0.018 to 0.048‰, whereas experiments exhibiting approximately 60–70% Zn adsorption yielded offsets ranging from approximately 0.004 to 0.09‰ (Table S2).

Among the native mosses, *S. denticulatum* generally exhibited relatively small isotopic offsets. Most values for *Sphagnum* were ≤0.05‰, despite substantial variations in pH and in the proportion of $Zn^{2+}$ adsorbed. The magnitude and direction of the fractionation were therefore broadly comparable among the different moss species, with no consistent species-specific enrichment pattern evident from the experimental dataset.

Zn isotope compositions were also determined for the in vitro-cultivated *S. palustre* clones-2a and 12a. Across the investigated adsorption range, the isotopic compositions measured for the cloned material overlapped with those obtained for native *Sphagnum* within analytical uncertainty (Fig. 3b). No systematic difference between the two clonal lines was apparent. Thus, under the experimental conditions investigated here, cultivation history did not produce a measurable systematic shift in the Zn isotope signature relative to field-grown *Sphagnum*.

**Table S3.** XANES linear-combination fitting results for Zn coordination environments in native and *in vitro*-cultivated moss samples.

| **Type of study** | **Sample** | **Species** | | | | | **R ($10^{-4}$)** | **$\chi^2$ ($10^{-5}$)** |
|---|---|---|---|---|---|---|---|---|
| | | | **ZnS** | **Cellulose** | **Acetate** | **Phosphate** | | |
| **Long-term** | Zn-2 | *Sphangum sp.* | 2 ± 1 % | 93 ± 2 % | 5 ± 3 % | | **11** | **26** |
| | Zn-4 | *Sphangum sp.* | | 100 ± 5% | | | **10** | **40** |
| | Zn-17 | *Sphangum sp.* | | 93 ± 2% | 7 ± 2 % | | **15** | **35** |
| | Zn-20 | *Sphangum sp.* | 1 ± 1 % | 92 ± 2% | 7 ± 8 % | | **21** | **53** |
| **Langmuir** | Zn-21 | *Sphangum sp.* | 3 ± 1 % | 81 ± 2% | 12 ± 2 % | 4 ± 5 % | **17** | **50** |
| | Zn-22 | *P. purum* | | 54 ± 2% | 6± 2 % | 40 ± 8 % | **7** | **20** |
| | Zn-23 | *Hypnum sp.* | 2 ± 2 % | 75 ± 2% | 11 ± 2 % | 12 ± 5 % | **16** | **46** |
| | Zn-24 | *P. purum* | | 87 ± 1% | 7 ± 2 % | 6 ± 3 % | **5** | **15** |
| **pH-edge** | Zn-25 | *Hypnum sp.* | | 97 ± 3 % | 3 ± 3 % | | **150** | **420** |
| | Zn-26 | *Sphangum sp.* | | 99 ± 6 % | 1 ± 3 % | | **160** | **460** |
| | Zn-27 | *P. purum* | | 100 ± 5% | 0 ± 5 % | | **150** | **450** |
| | Zn-28 | *B. rutabulum* | | 100 ± 5% | 0 ± 3 % | | **140** | **400** |
| | Zn-29 | *Hypnum sp.* | | 100 ± 5% | | 0 ± 3 % | **47** | **125** |
| | Zn-30 | *Sphangum sp.* | | 100 ± 5% | 0 ± 2 % | | **20** | **50** |
| | Zn-31 | *P. purum* | | 100 ± 5% | | 0 ± 3 % | **50** | **130** |
| | Zn-32 | *B. rutabulum* | | 100 ± 5% | 0 ± 2 % | | **12** | **30** |
| | Zn-37 | *Hypnum sp.* | | 91 ± 2 % | 9 ± 3 % | | **85** | **210** |
| | Zn-38 | *Sphangum sp.* | | 94 ± 2 % | 6 ± 3 % | | **95** | **250** |
| | Zn-39 | *P. purum* | | 94 ± 2 % | 6 ± 2 % | | **38** | **100** |
| | Zn-33 | *Hypnum sp.* | | 98 ± 1 % | 2 ± 8 % | | **20** | **50** |
| | Zn-35 | *P. purum* | | 96 ± 1 % | 4 ± 1 % | | **10** | **25** |
| | Zn-42 | *Sphagnum* clone C2a2 | | 95 ± 2% | 2 ± 1 % | 3 ± 1 % | **7** | **17** |
| **Growth clones** | Zn-47 | *Sphagnum* clone C2a2 | | 47 ± 4% | | 52 ± 10 % | **35** | **90** |
| | Zn-49 | *Sphagnum* clone C2a2 | | 54 ± 3% | | 46 ± 5 % | **9** | **22** |
| | Zn-51 | *Sphagnum* clone C2a2 | | 77 ± 4% | | 23 ± 4 % | **40** | **85** |

| | | | | | | | | |
|---|---|---|---|---|---|---|---|---|
| **Intact Biomass** | Zn-52 | *Sphangum sp.* | | 100 ± 2% | | 4 ± 5 % | **100** | **220** |
| | Zn-53 | *P. purum* | | 100 ± 4% | | 4 ± 5 % | **78** | **180** |
| | Zn-54 | *Hypnum sp.* | | 77 ± 6% | | 23 ± 15 % | **95** | **240** |
| | Zn-55 | *P. purum* | | 80 ± 5% | | 20 ± 10 % | **72** | **170** |

### Text to Table S3 — XANES contributions by experimental series

In the long-term *Sphagnum* experiments, cellulose accounted for approximately 92–100% of the fitted spectra, with acetate contributions of up to about 7%. Nominal ZnS contributions of approximately 1–2% were returned for some samples.

Greater compositional variability was observed in the adsorption-isotherm samples. Cellulose-like contributions ranged from approximately 54 to 87%, whereas phosphate-associated Zn reached up to 40% in one sample. Acetate generally represented a minor component, ranging from approximately 6 to 12%, and nominal ZnS contributions of 2–3% were obtained for some spectra.

The pH-edge series showed a substantially more homogeneous XANES signature. Cellulose-like coordination accounted for approximately 91–100% of the fitted spectra, while acetate or phosphate contributions were generally ≤9%. The *S. palustre* clone C-2a sample analyzed within this series similarly yielded 95 ± 2% cellulose-like coordination, together with 2 ± 1% acetate and 3 ± 1% phosphate.

A contrasting pattern was observed in the in vitro-grown clonal samples of the growth series (Zn-47 to Zn-51). In Zn-47, Zn-49, and Zn-51, cellulose-like contributions increased from 47 ± 4 to 77 ± 4%, whereas the phosphate-associated fraction decreased from 52 ± 10 to 23 ± 4%. These samples therefore contained some of the largest phosphate contributions observed in the dataset. The intact-biomass series (Zn-52 to Zn-55) also showed variable XANES compositions: some spectra were reproduced almost entirely by cellulose-like coordination, whereas others required phosphate contributions of approximately 20–23%.

The largest phosphate fractions occurred in selected clonal and 41-series samples, reaching approximately 40–52% of the fitted Zn signal. In contrast, the long-term and most pH-edge samples were dominated by cellulose-like coordination.

Within the practical resolution of the XANES linear-combination analysis, no significant contributions from metallic Zn, ZnO, $ZnCO_3$, Zn citrate, or $Zn_2SiO_4$ were required to reproduce the moss spectra. Small nominal ZnS fractions, generally below 5%, were obtained for a limited number of samples. Because these contributions are below the approximate resolution of the XANES fitting, they should be considered indicative rather than quantitatively robust.

**Table S4.** EXAFS-derived first-shell structural parameters for Zn in native and *in vitro*-cultivated moss samples. N is the resulting first-shell coordination numbers, R is the interatomic distances, and $\sigma^2$ is the Debye–Waller factors ($\sigma^2$). The "r" or goodness of fit of the curve to the original data.

| Type of study | Sample | | Scatter atom | R | N | $\sigma^2$ | r |
|---|---|---|---|---|---|---|---|
| **Long-term** | Zn-2 | *Sphangum sp.* | Zn-O | 2.029 | 4.8 | 0.0033 | 0.0100 |
| | Zn-4 | *Sphangum sp.* | Zn-O | 2.136 | 0.4 | 0.0040 | 0.020 |
| | | | Zn-S | 2.011 | 5.2 | 0.0068 | |
| | Zn-17 | *Sphangum sp.* | Zn-O | 2.025 | 5.6 | 0.0075 | 0.011 |
| | Zn-20 | *Sphangum sp.* | Zn-O | 2.033 | 6.0 | 0.0044 | 0.015 |
| | | | Zn-S | 2.079 | 0.4 | 0.0055 | |
| **Langmuir** | Zn-21 | *Hypnum sp.* | Zn-O | 2.049 | 6.0 | 0.0080 | 0.008 |
| | | | Zn-S | 2.134 | 0.6 | 0.0050 | |
| | Zn-22 | *Sphangum sp.* | Zn-O | 2.027 | 4.8 | 0.0074 | 0.010 |
| | Zn-23 | *P. purum* | Zn-O | 2.044 | 6.0 | 0.0065 | 0.009 |
| | | | Zn-S | 2.085 | 0.8 | 0.0050 | |
| | Zn-24 | *B. rutabulum* | Zn-O | 2.042 | 4.16 | 0.0035 | 0.012 |
| **pH-edge** | Zn-25 | *Hypnum sp.* | Zn-O | 2.040 | 6.8 | 0.0070 | 0.004 0.008 |
| | Zn-26 | *Sphangum sp.* | Zn-O | 2.035 | 6.4 | 0.0063 | 0.008 |
| | Zn-27 | *P. purum* | Zn-O | 2.040 | 7.6 | 0.0046 | 0.015 |
| | Zn-28 | *B. rutabulum* | Zn-O | 2.027 | 6.4 | 0.0074 | 0.021 |
| | Zn-29 | *Hypnum sp.* | Zn-O | 2.007 | 6.8 | 0.0081 | 0.01 |
| | Zn-30 | *Sphangum sp.* | Zn-O | 2.001 | 6.4 | 0.0080 | 0.013 |
| | Zn-31 | *P. purum* | Zn-O | 2.018 | 5.2 | 0.0060 | 0.016 |
| | Zn-32 | *B. rutabulum* | Zn-O | 1.996 | 6.8 | 0.0090 | 0.009 |
| | Zn-37 | *Hypnum sp.* | Zn-O | 2.033 | 6.8 | 0.0074 | 0.003 |
| | Zn-38 | *Sphangum sp.* | Zn-O | 2.044 | 7.6 | 0.0101 | 0.006 |
| | Zn-39 | *P. purum* | Zn-O | 2.014 | 7.3 | 0.0095 | 0.014 |
| | Zn-33 | *Hypnum sp.* | Zn-O | 2.016 | 6.0 | 0.0084 | 0.020 |
| | Zn-35 | *P. purum* | Zn-O | 2.016 | 6.4 | 0.0084 | 0.008 |
| | Zn-42 | *Sphagnum* clone C2a2 | Zn-O | 1.981 | 6.0 | 0.0077 | 0.02 |
| **Growth clones** | Zn-47 | *Sphagnum* clone C2a2 | Zn-O | 1.963 | 3.6 | 0.0039 | 0.031 |

| | | | | | | | |
|---|---|---|---|---|---|---|---|
| | Zn-49 | *Sphagnum* clone C2a2 | Zn-O | 2.018 | 4.0 | 0.0035 | 0.030 |
| | Zn-51 | *Sphagnum* clone C2a2 | Zn-O | 2.029 | 4.4 | 0.0048 | 0.006 |
| **Intact Biomass** | Zn-52 | *Sphangum sp.* | Zn-O | 2.005 | 4.9 | 0.0045 | 0.030 |
| | Zn-53 | *P. purum* | Zn-O | 2.035 | 4.0 | 0.0034 | 0.06 |
| | Zn-54 | *Hypnum sp.* | Zn-O | 1.974 | 3.6 | 0.0040 | 0.07 |
| | Zn-55 | *P. purum* | Zn-O | 1.961 | 4.8 | 0.0041 | 0.03 |

### Text to Table S4 — Detailed EXAFS structural parameters and isotope–structure comparisons

Debye–Waller factors were generally between approximately 0.003 and 0.010 Å² (Table S4), indicating substantial variability in the average local Zn coordination environment among the different moss materials and experimental conditions.

The pH-edge series exhibited the highest fitted coordination numbers. For these samples, coordination numbers generally ranged from 5.2 to 7.6, with Zn–O distances of approximately 2.00–2.04 Å. In contrast, the in vitro-grown samples (Zn-47 to Zn-51) showed lower coordination numbers, ranging from 3.6 to 4.4, with Zn–O distances between approximately 1.96 and 2.03 Å. The intact-biomass series (Zn-52 to Zn-55) similarly displayed relatively low coordination numbers (4 to 5) and Zn–O distances between approximately 1.96 and 2.04 Å. The C2a clone sample showed an intermediate-to-high coordination number (6.0) at a shorter Zn–O distance of 1.981 Å.

No simple monotonic relationship between Zn–O distance and coordination number was observed across the complete dataset. Nevertheless, samples characterized by the lowest coordination numbers, particularly those from the growth samples (Zn-47 to Zn-51) and intact reference samples (Zn-52 to Zn-55), tended to exhibit shorter average Zn–O distances (approximately 1.96–2.03 Å) than samples with the highest fitted coordination numbers in the pH-edge series (approximately 2.00–2.04 Å).

For a limited number of samples, incorporation of an additional Zn–S scattering path improved the EXAFS description. Zn–S contributions were included for H-41, P-41, Zn-4, and Zn-20 (Table S4). In H-41, P-41, and Zn-20, the presence of a Zn–S contribution is consistent with the small nominal ZnS fractions independently returned by the XANES linear-combination analysis. Because these sulfur-associated fractions are minor and close to or below the practical sensitivity of the XANES fitting, the Zn–S contribution should be considered evidence for a minor sulfur-containing coordination environment rather than a quantitatively dominant Zn species.

Overall, the EXAFS results showed that Zn retained by the mosses is predominantly coordinated by O-containing ligands, but that the local first-shell environment varied substantially among samples in both coordination number and Zn–O distance.

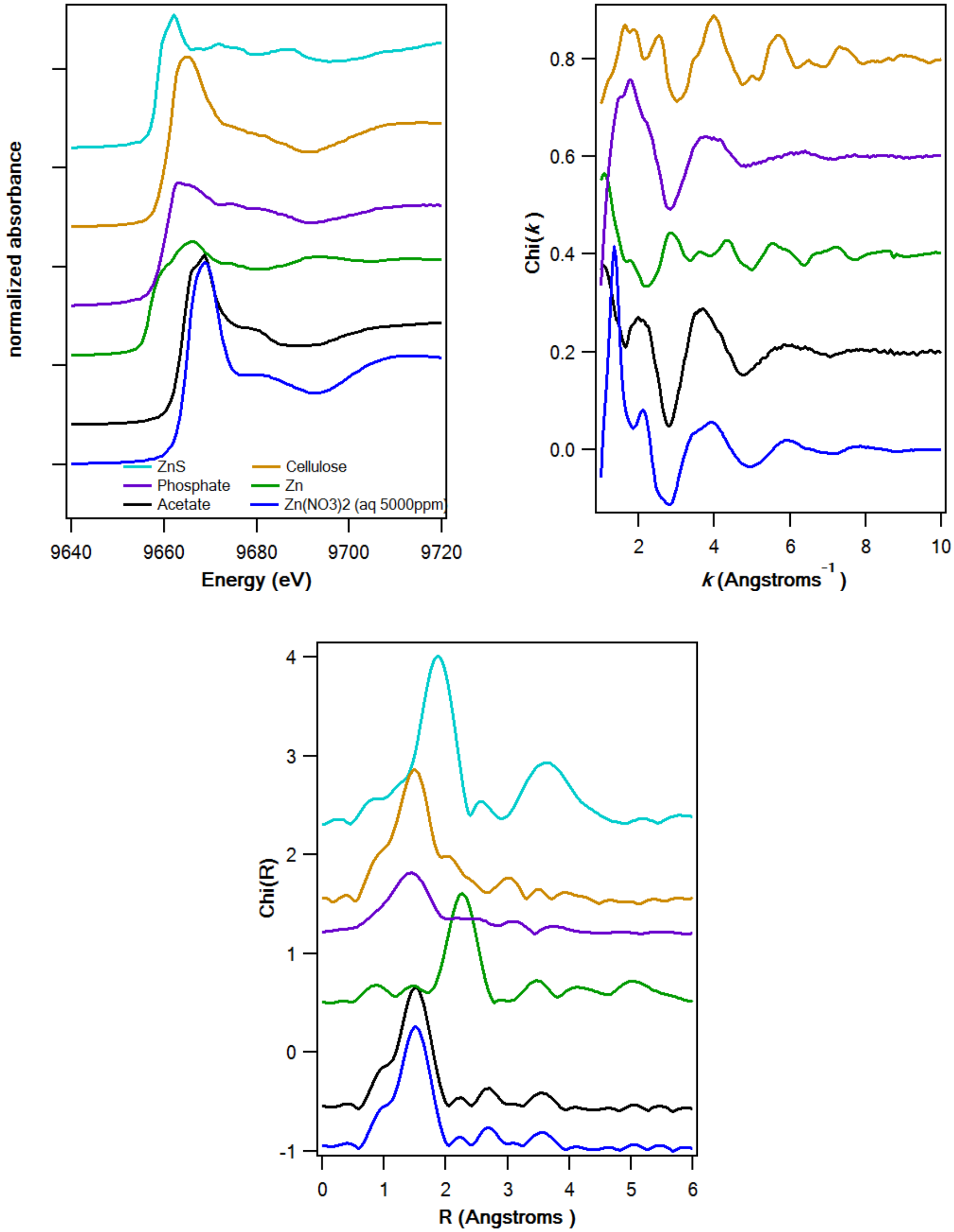

**Figure S1.** Zn K-edge XAS spectra of selected Zn reference compounds used for spectroscopic interpretation. (a) Normalized XANES spectra, (b) EXAFS χ(k) functions, and (c) corresponding Fourier transforms for ZnS, Zn phosphate, Zn acetate, Zn-associated cellulose, metallic Zn, and aqueous $Zn^{2+}$ prepared from $Zn(NO_3)_2$.

The reference compounds exhibited distinct near-edge features and edge positions, providing a set of spectral end-members for evaluating the local coordination environments of Zn in the moss samples.

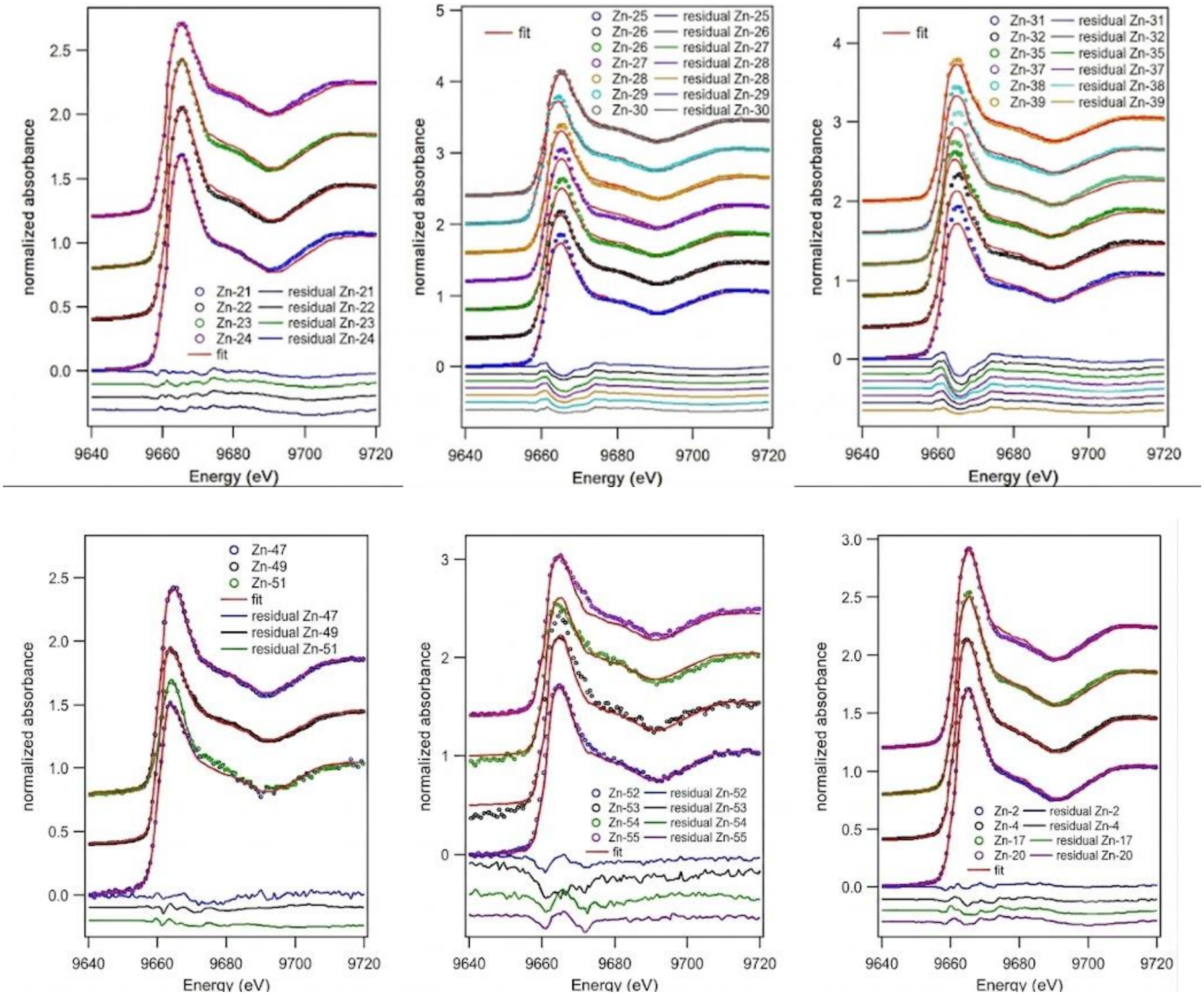

**Figure S2.** Zn K-edge XANES spectra and linear-combination fits for Zn associated with native and *in vitro*-cultivated moss samples. The panels include samples from the pH-edge experiments (XAS series), adsorption-isotherm experiments, *in vitro*-grown clonal material (growth series), intact biomass (Reference), and long-term *Sphagnum* adsorption experiments. Experimental spectra are shown together with the best-fit linear combinations, and residuals are displayed below each spectrum. Spectra are vertically offset for clarity. The relative contributions of the fitted Zn coordination environments are reported in **Table S3.** Although the overall spectral shapes were generally well reproduced by the linear-combination fits (Fig. S2), systematic residuals remained for some samples, particularly around the intensity of the main absorption-edge peak. The edge position and overall spectral envelope were nevertheless reproduced more closely by cellulose- and phosphate-containing references than by the other reference compounds. These residual differences indicate that the measured moss spectra cannot in all cases be represented perfectly as simple linear combinations of the available reference spectra and may reflect variations in the local geometry of Zn that are not fully captured by the selected standards.

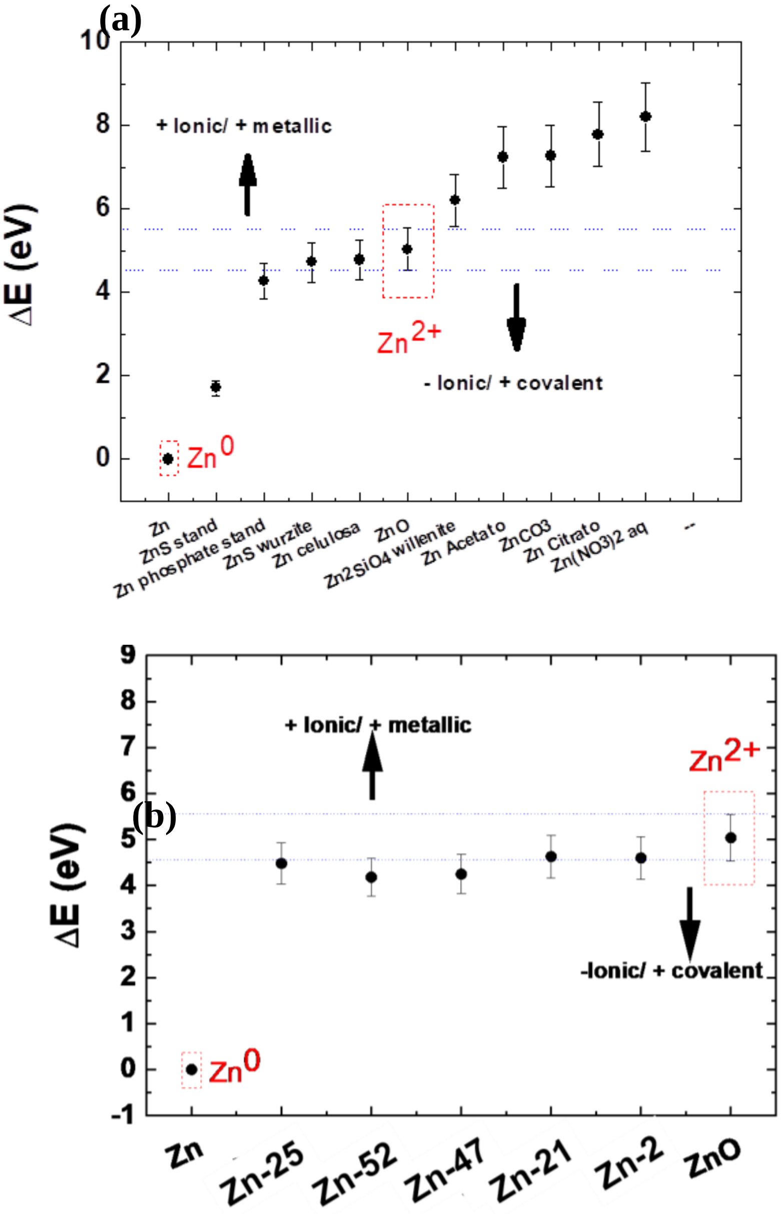

**Figure S3.** Zn K-edge absorption-edge shifts relative to metallic Zn for reference compounds and moss-associated Zn. (a) Relative edge-energy shifts of Zn reference compounds representing different inorganic, organic, and aqueous coordination environments. (b) Relative edge-energy shifts for the different moss experimental series compared with metallic Zn and aqueous $Zn^{2+}$. Moss-associated Zn exhibits a comparatively narrow range of positive edge shifts, whereas the reference compounds span a wider range of ΔE values. All moss samples displayed positive shifts relative to Zn, with mean values clustering within a comparatively narrow interval of approximately 4–5 eV. In contrast, the reference compounds covered a substantially wider range of edge shifts. The aqueous Zn reference occurred at higher ΔE than the moss samples, whereas several solid and organically coordinated Zn references displayed edge positions closer to those observed for the moss-associated Zn.

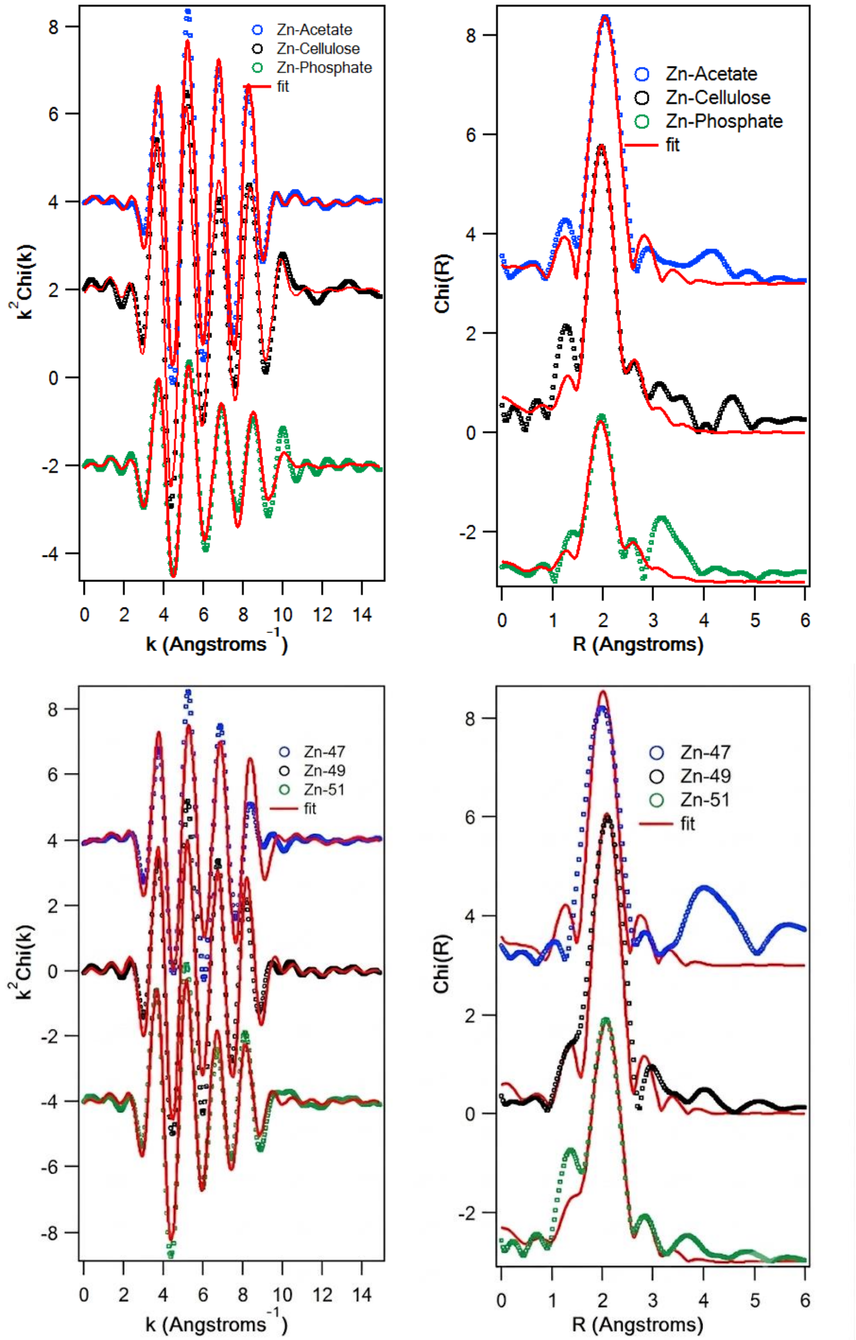

**Figure S4.** Zn K-edge EXAFS spectra and corresponding Fourier transforms for Zn associated with native and *in vitro*-cultivated moss samples.

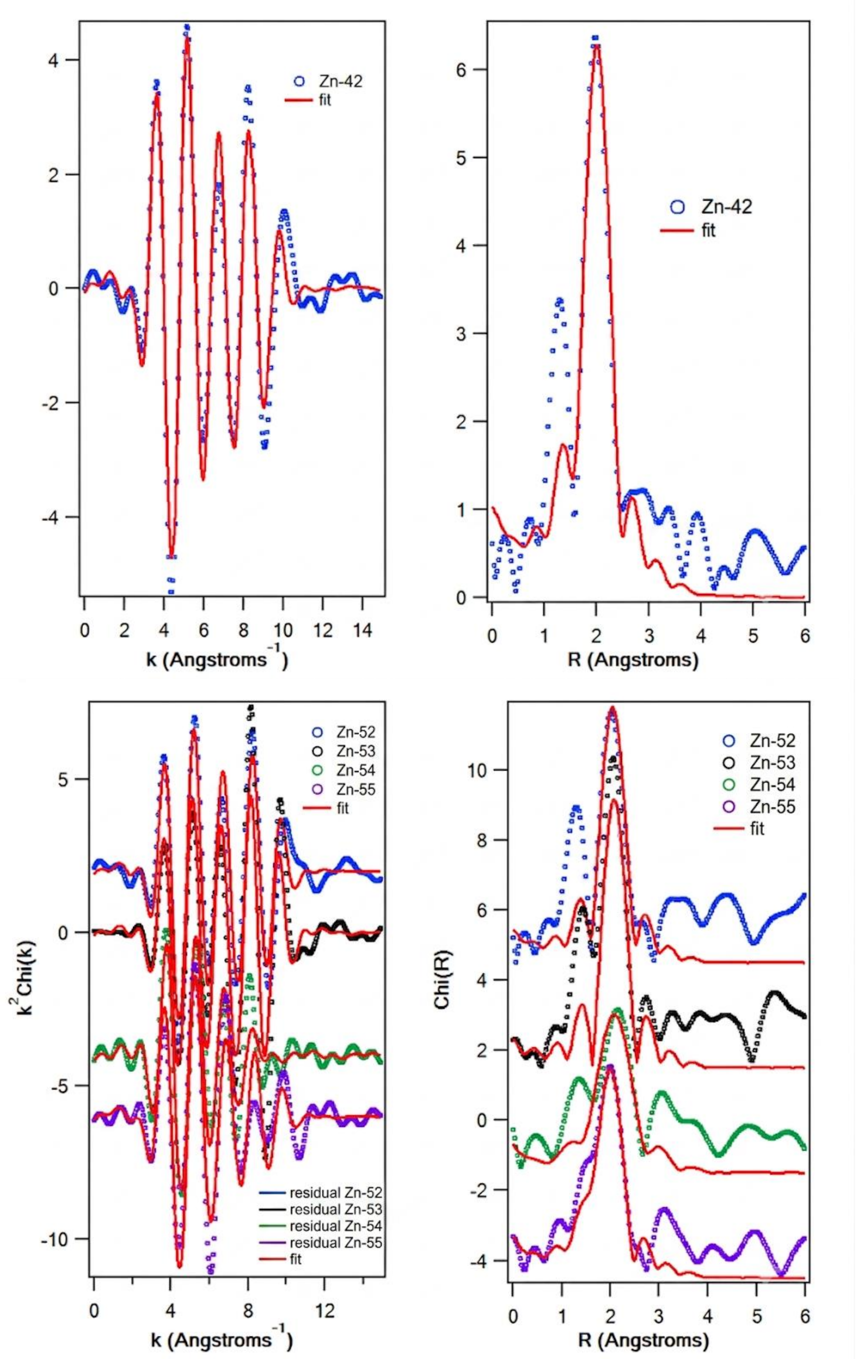

**Figure S4.** Zn K-edge EXAFS spectra and corresponding Fourier transforms for Zn associated with native and *in vitro*-cultivated moss samples.

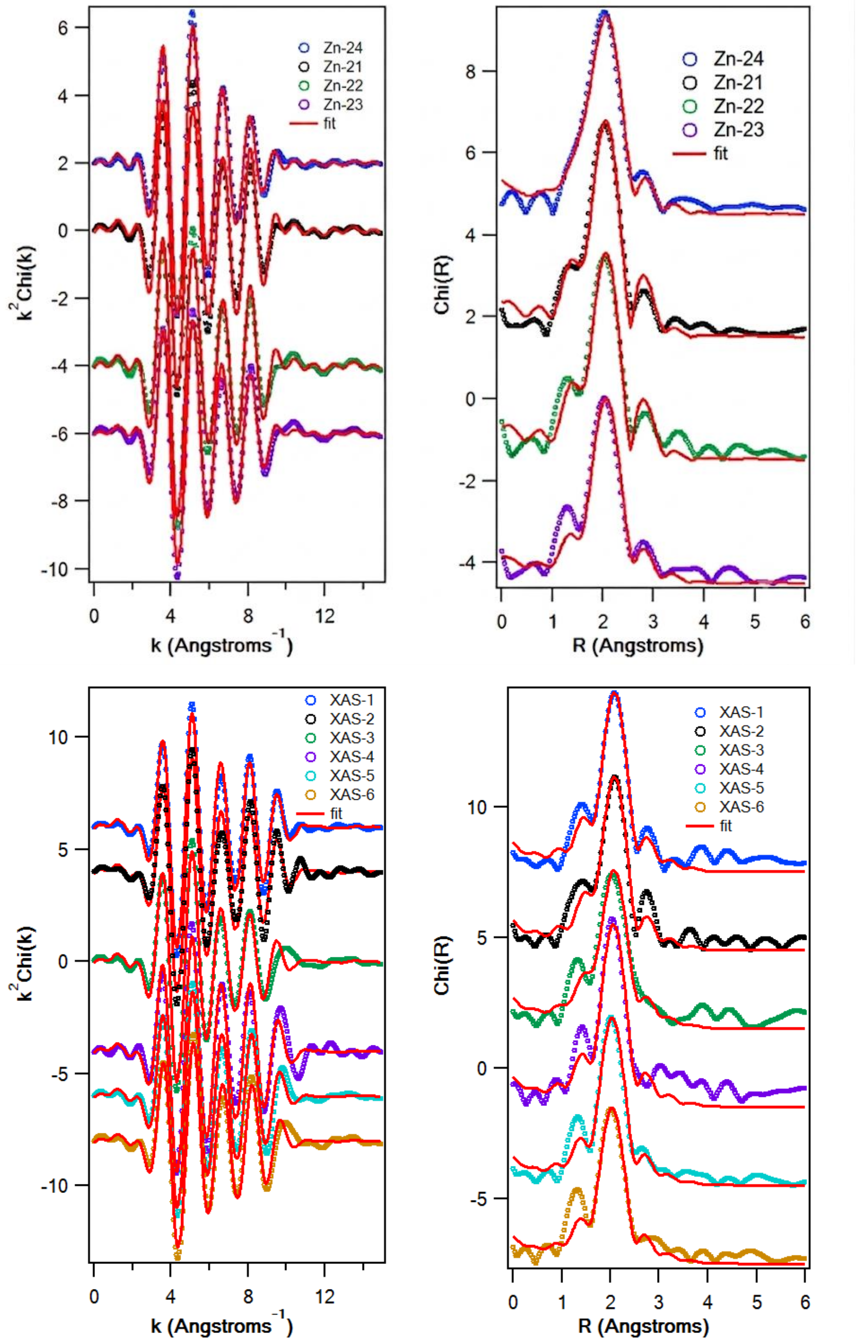

**Figure S4.** Zn K-edge EXAFS spectra and corresponding Fourier transforms for Zn associated with native and *in vitro*-cultivated moss samples.

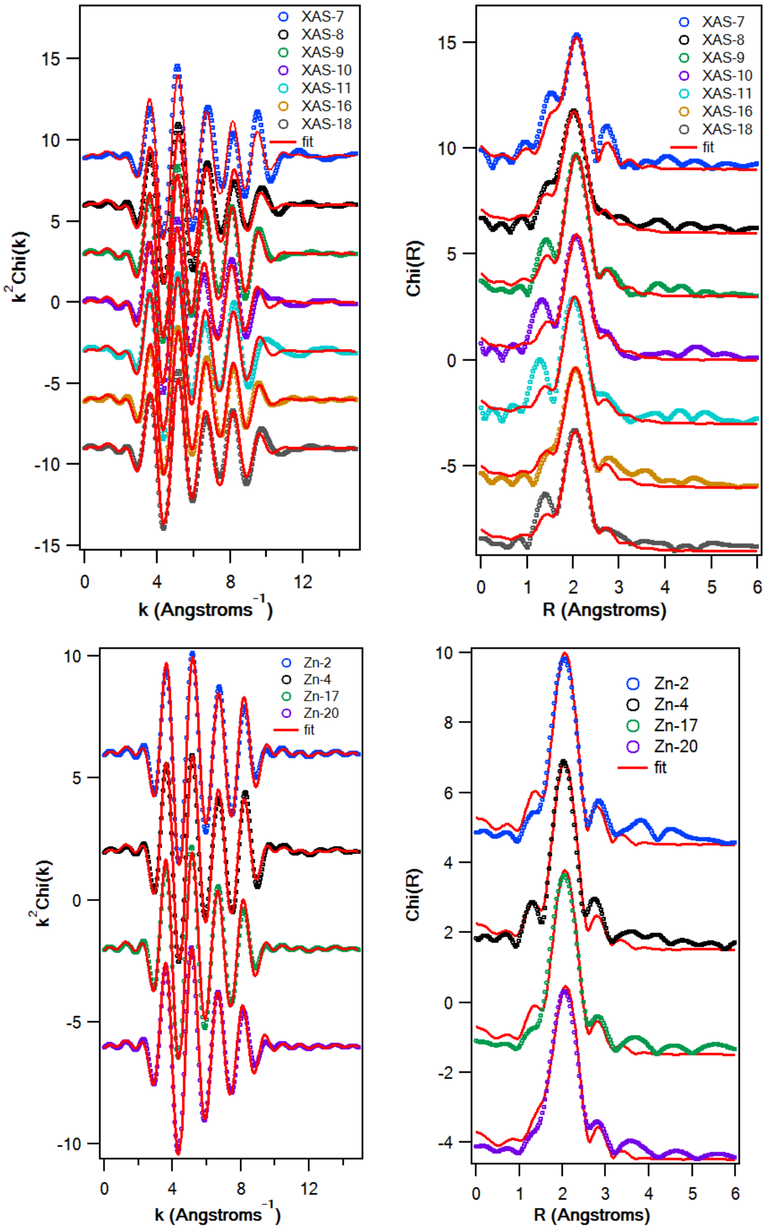

**Figure S4.** Zn K-edge EXAFS spectra and corresponding Fourier transforms for Zn associated with native and *in vitro*-cultivated moss samples. Experimental $k^2\chi(k)$ signals are shown together with the corresponding best-fit models, and their Fourier transforms are presented in R-space. The different panels include Zn reference environments and samples from the *in vitro*-grown clonal series (growth, C2a and C12-a), intact-biomass series (R), adsorption-isotherm experiments, pH-edge experiments, and long-term *Sphagnum* adsorption experiments. Spectra are vertically offset for clarity. EXAFS fits were based predominantly on Zn–O scattering paths, with an additional Zn–S contribution included where required. The resulting first-shell coordination numbers (N), interatomic distances (R), and Debye–Waller factors ($\sigma^2$) are reported in Table S4. The experimental signals were generally well reproduced by the fitted models over the measured range, and the Fourier transforms were dominated by a first-shell contribution centered at approximately 2 Å.
